# The Pan-European Impact of the Balkan Hunter-Gatherers

**DOI:** 10.64898/2026.08.05.743150

**Authors:** Daniel Tabin, Dušan Borić, Arev P. Sümer, Gregory Soos, Francesca Alhaique, Clive Bonsall, Adina Boroneant, Francesca Candilio, Emanuela Cristiani, Tracy Cullen, Ivana Fiore, Charoula M. Fotiadou, Stella Katsarou, Adisa Lepić, Ana Marić, Alana Masciana, Rita T. Melis, Margherita Mussi, Anastasia Papathanasiou, Catherine Perlès, T. Douglas Price, Kristine Korzow, Adamantios Sampson, Andrei Soficaru, Antonio Tagliacozzo, Kevin Uno, Orhan Efe Yavuz, Swapan Mallick, Iosif Lazaridis, Nick Patterson, Cosimo Posth, Ron Pinhasi, Nadin Rohland, Mateja Hajdinjak, Esther Brielle, David Reich

**Affiliations:** Department of Human Evolutionary Biology, Harvard University, Cambridge, MA 02138, USA; Department of Environmental Biology, Sapienza University of Rome, Rome, Italy; Department of Anthropology, New York University, New York, NY 10003, USA; The Italian Academy for Advanced Studies in America, Columbia University, New York, NY 10027, USA; Department of Evolutionary Genetics, Max Planck Institute for Evolutionary Anthropology, Leipzig, Germany; Department of Genetics, Harvard Medical School, Boston, MA 02115, USA; Howard Hughes Medical Institute, Harvard Medical School, Boston, MA 02115, USA; Museo delle Civiltà, 00144 Rome, Italy; School of History, Classics and Archaeology, University of Edinburgh, UK; “Vasile Pârvan” Institute of Archaeology, Romanian Academy, Bucharest, Romania; DANTE – Diet and Ancient Technology Lab, Department of Odontostomatological and Maxillofacial Sciences, Sapienza University of Rome, Rome, Italy; American School of Classical Studies at Athens, Athens, Greece; Archaeo- and Palaeogenetics, Institute for Archaeological Sciences, Department of Geosciences, University of Tübingen, Tübingen 72074, Germany; HUMAN ORIGINS—Cluster of Excellence for Integrative Human Origins Studies (EXC 3101), University of Tübingen, Tübingen 72074, Germany; Ephorate of Palaeoanthropology and Speleology, Hellenic Ministry of Culture, Athens 11636, Greece; National Museum of Bosnia and Herzegovina, Zmaja od Bosne, Sarajevo, Bosnia and Herzegovina; Lamont-Doherty Earth Observatory of Columbia University, Palisades, NY 10964, USA; Dipartimento Scienze Chimiche e Geologiche, Università degli Studi di Cagliari Cittadella Universitaria, Blocco A,09042 Monserrato, Italy; ISMEO-The International Association for Mediterranean and Oriental Studies, Corso Vittorio Emanuele II 244, 00186 Roma, Italy; Université Paris Nanterre, CNRS, UMR 8068, France; Department of Anthropology, University of Wisconsin-Madison, USA; Department of Anthropology, Texas A&M, TX 77843, USA; Department of Mediterranean Studies, University of the Aegean, Rhodes, Greece; Department of Paleoanthropology/Human Osteology, Institute of Anthropology, Bucharest 050711, Romania; 1 Decembrie 1918 University of Alba Iulia, Faculty of History, Letters and Educational Sciences, Alba Iulia 510009, Romania; Senckenberg Centre for Human Evolution and Palaeoenvironment at the University of Tübingen, Tübingen 72074, Germany; Broad Institute of MIT and Harvard, Cambridge, MA 02142, USA; Department of Evolutionary Anthropology, University of Vienna, A-1030 Wien, Austria; CNR-IGAG, Piazzale Aldo Moro 7, 00185 Rome, Italy; Department of Earth and Planetary Sciences, Harvard University, Cambridge, MA 02138, USA

## Abstract

During the Last Glacial Maximum (LGM) 26–19 thousand years ago (kya), Europe had three principal refugia not covered by ice: the Iberian, Italian, and Balkan peninsulas^1,2^. While the Iberian and Italian refugia have been studied with ancient DNA^3–7^, the legacy of the Balkan refugium has remained a mystery due to a lack of genetic data 30**–**12 kya. We present genome-wide data from nine newly reported individuals: three Epipaleolithic from Romania, one Epipaleolithic from Bosnia and Herzegovina, three Mesolithic from Greece, one Epipaleolithic from Southern Italy, and one Mesolithic from Sardinia. We find that the Epipaleolithic individuals from Romania and the Mesolithic individual from mainland Greece were from a previously unsampled population that was the primary source for later European hunter-gatherers. Westward expansion of Balkan Epipaleolithic people into the Italian Peninsula and mixture with a minority contribution from pre-LGM Italians, produced a population that then spread further west to become the primary ancestry of western European Mesolithic people. Northeastward expansion, bypassing Italy, contributed the European ancestry source of Scandinavian and Eastern European hunter-gatherers. Southeastward expansion to the Aegean led to Anatolian-European mixtures before the spread of farming.

---

Humans survived in the Balkan refugium during the LGM^8^ at a time when northern Europe was largely uninhabitable. But in contrast to the other two southern European refugia^3^—Iberia and Italy—the only reported ancient DNA from post-LGM hunter-gatherers in this region comes from individuals who lived at least eight millennia after the ice began to recede. The only substantial published data from Balkan hunter-gatherers comes from people living in the Iron Gates area of the Danube (present-day Romania and Serbia),^7^ dating to 11–8 kya, whose ancestry was broadly similar to other Mesolithic groups from Western Europe^3^. It has been a mystery whether their ancestors originated in the Italian or Balkan refugia, as the only clear genetic difference with the Mesolithic hunter-gatherers of Italy was that the Iron Gates individuals^3^ had additional affinity to Siberian Paleolithic populations. Were the Balkans a mixing ground of people from the well-sampled Italian and Eastern European hunter-gatherer populations with little contribution of the local refugium? Or is the WHG+EHG model an artefact of poorer sampling in the Balkans, underestimating the contribution of the Balkan refugium? Archaeology on both sides of the Adriatic also points to a shared Epigravettian culture between Italy and the Balkans, making it difficult to infer deeper origins from the archaeology alone^9–11^. Moreover, archaeological research is unevenly distributed across the two sides of the Adriatic, with the eastern Adriatic remaining substantially less investigated than Italy and other regions of western Europe. This imbalance hinders our ability to characterize the direction, timing and origins of material culture change during the transition from the Gravettian to the Early Epigravettian (c. 26–25 kya) and from the Early to the Late Epigravettian (c. 17.6–17.1 kya). These cultural transformations and associated shifts in mobility and land-use strategies, unfolded against a backdrop of major climatic and environmental changes, including a global sea-level fall of up to 120–130 m during the LGM that exposed a vast land bridge known as the Great Adriatic-Po Region connecting the Italian and Balkan peninsulas^12^.

The nomenclature used for the genetic ancestry of hunter-gatherer populations in Europe has changed across publications^3,5,13^. Since our goal is to compare the ancestry impact of the people of the three southern refugia, for clarity, we use a terminology based on archaeological culture and region^9^. We use “Epipaleolithic Spanish” (Epi-Spanish) to refer to groups commonly denoted in the genetic literature as the “Fournol” cluster^3^, “Epipaleolithic Italian” (Epi-Italian) to refer to groups previously denoted as the “Villabruna” cluster^3,5^, and “Epipaleolithic Balkan” (Epi-Balkan) to refer to the distinctive population whose ancestry we describe here (Table 1).

**Table 1:** Terminology. We use a consistent naming scheme based on the location and time this ancestry appears, sometimes further annotated with archaeological culture. We give other names by which these ancestries were known in the second column.

| Name of group | Previously used group names |
| --- | --- |
| Balkan Epipaleolithic (Epi-Balkan) | Not previously described |
| Italian Epipaleolithic (Epi-Italian) | Western Hunter-Gatherer (WHG)<sup>13</sup>, Villabruna Cluster<sup>13</sup> |
| Spanish Epipaleolithic (Epi-Spanish) | Fournol Cluster<sup>3</sup>, Magdalenians, Solutreans |
| Sardinian Mesolithic | Not previously described |
| Post-LGM Northwest European | WHG<sup>13</sup>, Oberkassel Cluster<sup>3</sup> |
| Central Eurasian Mesolithic (CEM) | Eastern Hunter-Gatherers (EHG)<sup>43</sup>, Sidelkino Cluster<sup>3</sup>, Forest Steppe Hunter-Gatherers (FSHG)<sup>18</sup>, Western Siberian Hunter-Gatherers (WSHG)<sup>38</sup> |
| French pre-LGM | Fournol Cluster<sup>3</sup>, French Gravettians<sup>3</sup>, Western Gravettians<sup>3</sup> |
| Spanish pre-LGM | Fournol Cluster<sup>3</sup>, Western Gravettians<sup>3</sup>, Spanish Gravettians<sup>3</sup> |
| Italian pre-LGM | Italian Gravettian<sup>3</sup>, Southern Gravettians<sup>3</sup>, Věstonice Cluster<sup>3</sup> |
| Central Europe pre-LGM | Věstonice Cluster<sup>5</sup>, Austrian Gravettian<sup>5</sup>, Czech Gravettian<sup>5</sup> |
| Paleolithic Siberians | Ancient North Eurasians (ANE)<sup>13</sup> |
| Anatolian Epipaleolithic | [unchanged]<sup>27</sup> |
| Anatolian Neolithic | [unchanged]<sup>48</sup> |
| Caucasus post-LGM | Caucasus Hunter-Gatherer (CHG)<sup>20</sup> |
| Caucasus pre-LGM | Dzudzuana Cluster<sup>49</sup> |
| Scandinavian Mesolithic | Scandinavian HG<sup>13</sup> |
| West Balkan Mesolithic | Croatian HG<sup>7</sup>, Balkan HG<sup>7</sup> |
| East Balkan Mesolithic | Serbian HG<sup>7</sup>, Balkan HG<sup>7</sup> |
| Iberian Mesolithic | Portuguese HG<sup>6</sup>, Spanish HG<sup>4</sup>, El Mirón Cluster<sup>5</sup> |
| Sicilian Mesolithic | Sicily HG<sup>25</sup> |

We prepared skeletal samples in dedicated clean rooms, successfully generated 3-33 double-^14,15^ or single-stranded^16^ libraries from each of nine individuals (a total of 100 libraries are newly reported here; Table S1), enriched them in-solution using reagents that target more than one million single nucleotide polymorphisms (SNPs)^17^, and sequenced the enriched products on Illumina instruments (Methods). The highest-quality data derives from three Romanian Epipaleolithic individuals from Climente II (14.5–11 kya), who we used in this study to represent the Balkan refugial population. We analyzed genetic data from these nine newly reported individuals along with that from 1,110 previously published ancient and modern samples (Table S2). The great majority of previously reported hunter-gatherers from Europe with genome-wide ancient DNA have been analyzed using in-solution enrichment^3–7,18–42^ (Table S1, S3). Due to the sparsity of comparative shotgun data from European hunter-gathers, shotgun sequencing and analyzing data from the single Balkan hunter-gatherer individual (from Climente II) for whom the proportion and amount of human data was high enough for non-enrichment based approaches to be economical, would not have added value to the fine-grained analyses of the population structure of European hunter-gatherers which are the focus of the present study.

We report three main findings. First, the Epi-Balkan population was closely related to, but genetically distinct from, the Epi-Italian population, reflecting an expansion of populations from the Balkans into the Italian Peninsula where they mixed in a previously undocumented way with a small proportion of ancestry from pre-LGM Italian Gravettian-associated populations. Second, Epi-Iberian, Epi-Italian, and Epi-Balkan ancestries differentially contributed to the formation of later European hunter-gatherers. Epi-Balkan ancestry contributed the bulk of Epipaleolithic and Mesolithic European ancestry: Epi-Italian ancestry (itself being primarily of Epi-Balkan origin) became common west of the Oder River, while Epi-Balkan ancestry contributed directly to nearly all of the refugial ancestry groups further east and in Scandinavia. Third, there were pre-Neolithic connections between Anatolia and the Balkans: individuals on an Aegean island in the Western Cyclades were a mixture of Anatolian and Epi-Balkan ancestries.

## Balkan people expanded southeastward mixing with Anatolians

The data from the Epipaleolithic individual from Badanj are important in showing that Epi-Balkan ancestry was present at least by 16 kya, while those from Franchthi reveal that such ancestry survived in an unadmixed form as recently as 10 kya. Outgroup-*f_3_* statistics show that the Romanian Epigravettians from Climente II, Franchthi, and Badanj individuals are similar (Table S3), which we confirmed using statistics of the form *f_4_*(Yoruba, X; Franchthi / Badanj, Romanian Epigravettian) for a variety of pre-LGM populations X. We computed this comparison with both the full and damage-restricted data for Franchthi and Badanj to remove contamination (Methods) (all |Z|<3; Table S2). In *qpWave*^43^, Badanj and Climente II cannot be rejected as descending from a single source of ancestry without mixture (p > 0.23) using a diverse set of reference populations; however, the model only marginally passes when Epi-Italians are moved into the outgroups (p ∼ 0.03), suggesting that the Balkan gene flow into Italy we document in what follows was closer to Badanj than to Climente II Epi-Balkan people. An alternative explanation is gene flow from Italy into the Balkans, which we can demonstrate did occur by at least 9 kya through our analysis of West Balkan Mesolithic individuals from Croatia.

Two individuals from the site of Maroulas on the Aegean island of Kythnos and dating to 8993–8651 cal BP and 8595–8431 cal BP, have a unique genetic profile intermediate between Balkan Epigravettians and Anatolians represented by the Epipaleolithic Anatolian from Pınarbaşı. Analysis using DATES^44^ confirms the Maroulas individuals have significant admixture between these two sources (Z>3.7), which is unexpected if this signal were an artifact of contamination. The sizes of segments of the two ancestries are what would be expected for mixture before the date of arrival of farming in the Balkans and Aegean islands: 1034±266 years (one standard error) before their radiocarbon dates (Figure 1B, Extended Data 2, 3, 4). In *qpAdm*, the Maroulas individuals exhibit far more Epi-Balkan-related ancestry than any Balkan Neolithic groups (∼30±6%) (Extended Data 4). Epi-Balkan ancestry is present even in the contaminated data from Maroulas 2, as models in which Epi-Balkan groups are moved into the right (see methods) are strongly rejected (p < 0.00001). Thus, before the onset of the Neolithic, there was gene flow between European hunter-gatherers and peoples of Anatolia. Further sampling would shed light whether this was a long-standing cline across the Aegean islands or whether this mixture was formed by population movements just before the start of the Neolithic.

**Figure 1:**
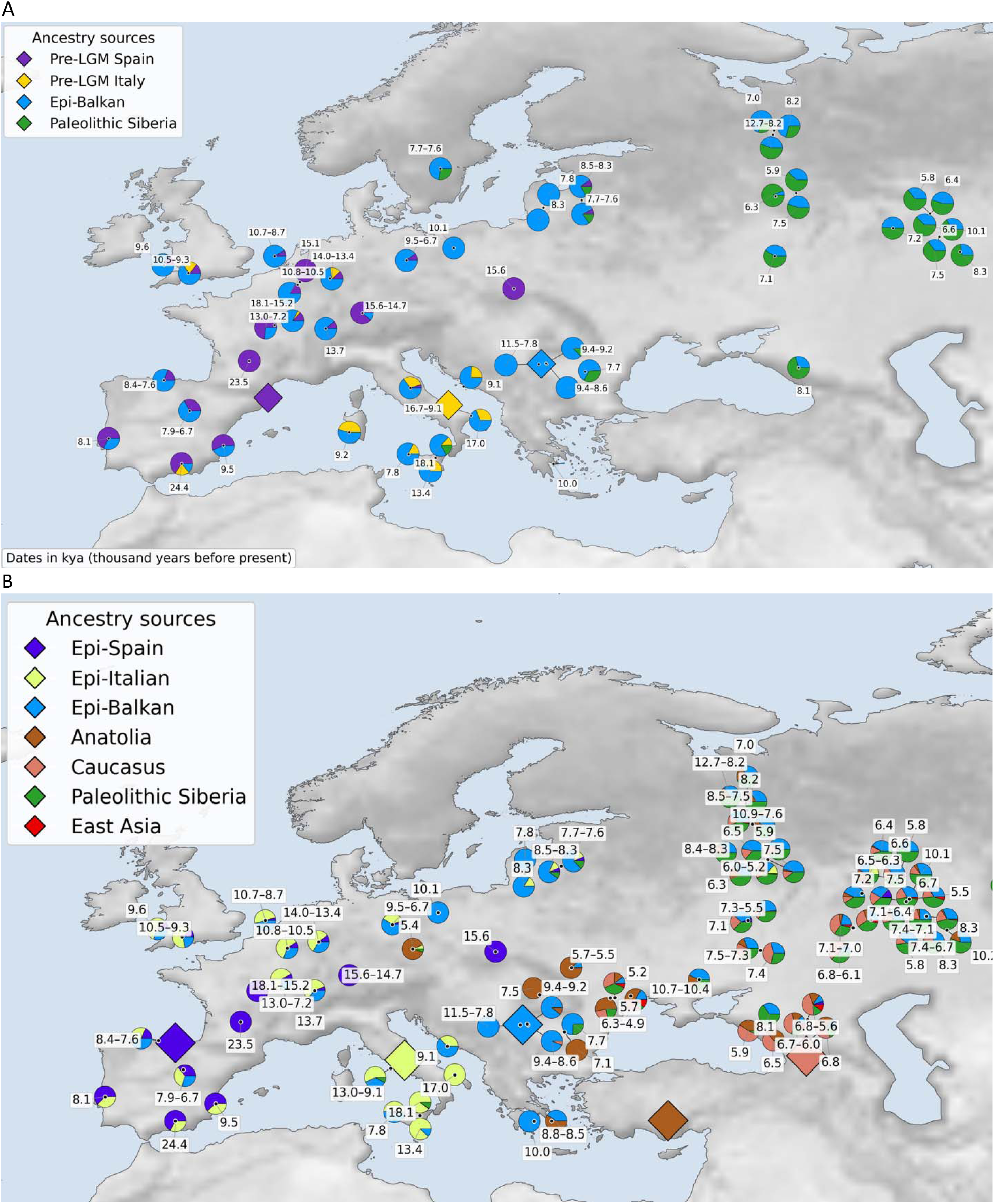
The ancestry of Europe’s hunter-gatherers after the Ice Age. (A) Modeling with proxies for the three refugial populations of southern Europe shows that the primary ancestry of post-LGM Europeans is from the Balkans. The simplest, passing, non-negative model with sources rotated to the right is plotted; hunter-gatherers not plotted require the additional sources in the second panel. (B) Modeling with post-LGM sources highlights how much of the Balkan hunter-gatherer ancestry spread via the Italian Peninsula in the West, but without any Italian contribution in the North and East. (See Methods and Tables S5-S9 for *qpAdm* modeling details and full results).

## Balkan people expanded westward via the Italian Peninsula

Despite a shared Epigravettian culture^9–11^, the Epi-Balkan and Epi-Italian populations are not a clade, which we confirmed by computing symmetry statistics of the form *f_4_*(Yoruba, X; Epi-Italian, Epi-Balkan) and observing many significant deviations for a range of older populations X (Table S2). For example, pre-LGM Gravettians from many parts of Europe, and especially pre-LGM individuals from Italy (Z-scores < -14), show excess affinity to Epi-Italians, while none do to Epi-Balkans (all Z-scores < 1.5). However, the two groups share far more genetic drift with each other than either do with preceding groups, indicating shared ancestry, as shown by symmetry statistics of the form *f_4_*(Yoruba, Epi-Italian; X, Epi-Balkan) and *f_4_*(Yoruba, Epi-Balkan; X, Epi-Italian) (Z-scores: 6.6–54.4, Table S4).

Motivated by these results, we tested whether Epi-Italian groups were consistent with being admixed between Balkan and pre-LGM Italian populations using *qpAdm*^43^. In the absence of Epipaleolithic Balkan genetic data, it was previously suggested that Epi-Italians derived from a Balkan migration that may have fully replaced the previous Italian Gravettian population^3^. Our work provides direct evidence that much of the ancestry of the Epi-Italian population indeed derives from Epi-Balkan people. However, in tension with the hypothesis of a full replacement, our sampling of Balkan people living after the LGM shows that the Epi-Italian population derives substantial ancestry from the pre-LGM Italian populations that preceded it. Many Italian hunter-gatherers (i.e. both the Epi-Italian population and later Mesolithic Italians) also require additional minor ancestry related to pre-LGM Spain to fit the data (Figure 1, Tables S5-S8). The best-fitting model shows that Epi-Italian individuals as a whole have ∼75% Epi-Balkan-related ancestry in addition to ∼20% Italian pre-LGM-related ancestry and ∼5% Spanish pre-LGM-related ancestry.

Epi-Balkan ancestry rose in Italy in post-glacial times and was geographically structured, consistent with an intrusive and spatially uneven process. The earliest (∼17.5 kya) Italian post-LGM individuals, from Apulia^45^, show lower proportions of Epi-Balkan ancestry (∼55%) than the slightly later (<17 kya) Mainland Epi-Italians (∼75%) (Figure 1). The point estimate of 55% Epi-Balkan ancestry in the newly reported isolated Sardinian Mesolithic individual suggests they may have preserved lower proportions of Balkan ancestry well into the Holocene. On the other hand, the Mesolithic population from Sicily (∼8 kya)^25^ shows one of the highest proportions of Epi-Balkan ancestry (∼80%), indicating continuing gene flow through the Mesolithic. Some ancestry differences are significant, such as *f_4_*(Mainland Italian Epigravettian, Early Sicilian Epigravettian; pre-LGM Italian, Epi-Balkan) being negative at |Z|>5 standard errors from zero. The evidence for higher Epi-Balkan ancestry in Sardinia than the mainland is not significant with *f_4_*(Mainland Italian Epigravettian, Mesolithic Sardinian; pre-LGM Italian, Epi-Balkan) |Z| < 2, which could reflect a limited power due to limited information from Sardinia despite our efforts to increase data quality (average 0.17-fold genomic coverage on targeted sites from pooling 24 ancient DNA libraries; Methods). The newly reported data from Romanelli in the south of Italy (Apulia) and dating to roughly 13 kya are indistinguishable with *qpAdm* from other mainland Italian Epigravettians.

Archaeologically, key questions remain regarding the timing, drivers and material correlates of the initial demic diffusion of Epi-Balkan ancestry into Italy. This process plausibly occurred during the LGM, possibly as early as 26 ka and certainly by c. 19–18 ka, when Epi-Balkan ancestry is already well documented in Italy. This interval coincides with the Early Epigravettian techno-complex and with the probable extensive occupation of the Great Adriatic-Po Region by both human and animal populations^12^. Subsequent glacier retreat after 17.5 ka and the onset of rapid sea-level rise after 16.5 ka likely triggered the contraction of populations from the Great Adriatic-Po Region and their dispersal into the mountainous regions surrounding the Adriatic Basin^46^. However, it remains unclear why and through which demographic mechanisms Epi-Balkan ancestry came largely to absorb Italian and other pre-LGM European ancestries.

Epi-Balkan ancestry reached Iberia too, but not in as high proportion as in Italy. As in Italy there was spatiotemporal variation in this mixture with proportions of Epi-Balkan ancestry increasing over time. *qpAdm* indicates that people from the Iberian Refugium during and just after the LGM derive around 85% of their ancestry from Iberian pre-LGM populations. The rest was a mixture of Italian pre-LGM and Epi-Balkan, although we cannot reject models in which Solutrean-related populations derive all of their ancestry from the Spanish pre-LGM, nor models in which Magdalenian-related populations have minor Epi-Balkan, but no pre-LGM Italian. As in the Italian Peninsula, later Iberian Mesolithic groups show increased (30–75%) Epi-Balkan ancestry (Table S3); however, the proportions in Spain remained lower than in Italy until the Neolithic.

In post-LGM Northwest Europe, Epi-Balkan people also made a major impact indirectly through their contribution to Epi-Italian people, who are the main proximal ancestry source, with additional contributions from Epi-Spanish (Figure 1B). This previously described mixture has been called the “Oberkassel cluster” after one of the earliest high-quality genomes with this signature^3^.

## Balkan people expanded north and east, bypassing the Italian Peninsula

While in Western Europe, much of the Balkan expansion shown in Figure 1 passed through Italy, incorporating Italian pre-LGM-related ancestry along the way, in Northeastern Europe and Central Asia, European refugial associated ancestry is consistent with being entirely Epi-Balkan in origin. Neither pre-LGM Spanish related nor pre-LGM Italian related ancestry is required. Models could not be rejected even when those sources are moved into the right populations for *qpAdm,* a stringent test that excludes a significant contribution from already admixed Epi-Italian or Epi-Spanish populations. This pattern differs from that of post-LGM Northwest European populations, which all require Italian pre-LGM ancestry, Spanish pre-LGM ancestry, or both (Tables S5-8).

To provide finer-grained insight into which parts of Mesolithic Europe were directly impacted by expansions from the Balkan refugium, and which were impacted by expansions from the Balkans via the Italian Peninsula, we developed a *qpAdm* model that replaced pre-LGM Iberians and Italians with the post-LGM populations of each region. We also added three additional populations from further east: Caucasus Hunter-Gatherers, Anatolian Hunter-Gatherers, and East Asians (Figure 1B, Extended Data 3, Table S5). These results confirm that in the West (from Northwest European to Iberian Mesolithic peoples), Epi-Balkan associated ancestry spread primarily via Italy. In contrast, a direct expansion from a Balkans Refugial-related population spread this ancestry northeastwards to Scandinavia, Northeastern Europe, and as far east as the Altai Mountains (Extended Data 3).

As previously reported^13,30,47^, ancestry related to Upper Paleolithic Siberians is found in Scandinavia and further east, with varying degrees of mixture from East to West, making Scandinavia part of a cline that extended through Eastern Europe and into Siberia^18^. Epi-Balkan related ancestry is definitively the western partner in this mixture process: across this whole range, there is no evidence of any Italian or Iberian influence, even in Scandinavia. Models that use an Epi-Italian source instead of an Epi-Balkan source fail (p << 0.000001). Models including both Epi-Balkan and Epi-Italian sources assign over 70% (±10%) of the ancestry to the Balkan source and negative ancestry to Italian (Table S5-S9).

## Conclusion

Across Eurasia, massive population turnovers occurred around the time of the Last Glacial Maximum, and groups that survived the LGM repopulated Eurasia rapidly, giving rise to the ancestors of present-day Eurasians. Here we show that the LGM population of the Balkans was the main source of Late Glacial and Mesolithic hunter-gatherer related ancestry across Europe and a major contributor to Central Eurasian Mesolithic hunter gatherer populations, while the populations of Iberian and Italian before the LGM largely went extinct. Despite further massive population turnover during the Neolithic, the Epi-Balkan population continues to influence the genetic ancestry of present-day people. The Epi-Balkan population was a major component of the ancestry of Eastern European Hunter-Gatherers (sometimes called the “Sidelkino cluster”), who in turn gave rise to the Yamnaya, who were around 16% Epi-Balkan. The Yamnaya, in turn, contributed ancestry not only to Europeans but also to Central and South Asians, highlighting the worldwide impact of Epi-Balkan people even today.

## Methods

### Terminology for population groups

A multiplicity of terms has been used in the archaeogenetic literature to refer to the hunter-gatherers of Europe. An early principled approach was to name groups after the oldest high-quality genome representing people of that cluster, following a philosophy of defining genetic relationships independent of archaeological data, enabling objective evaluation of both lines of evidence^3^. In the present study, our hypotheses are explicitly spatiotemporal, and so we use geography and time to label groups, following another established precedent in the ancient DNA literature^9^.

A list of individuals which constitute the populations used in various analyses can be found in Table S10.

#### Sampling ancient individuals

All skeletal remains used to generate data presented in this paper were sampled in ancient-DNA clean rooms at Harvard Medical School in the USA or the Max Planck Institute for Evolutionary Anthropology in Leipzig, Germany^39^.

#### Ancient-DNA data generation

We present genome-wide data for nine individuals: three from Mesolithic Greece (Franchthi Cave and Maroulas), three from Epipaleolithic Romania (Climente II), one from Mesolithic Sardinia (S’Omu e S’Orku), one from Epipaleolithic Herzegovina (Badanj), and one from Epipaleolithic Apulia, Italy (Romanelli). For all nine individuals, the data come from sequencing the products of in-solution enrichment of more than one million single-nucleotide polymorphisms (SNPs)^17,43,50^ derived from 3 to 33 double- or single-stranded libraries each (Table S1). We analyzed the data together with previously published data^3–7,18–42^.

The oldest uncontaminated newly reported individual is from the site of Climente II in Romania, dating to the Epigravettian period, and is radiocarbon-dated to 14842–14100 years cal BP (12687±45 BP; after a correction for the freshwater reservoir effect: 12349±63 BP, OxA-40804) (95.4% calibrated confidence interval). This individual was a male (genetic ID I5410, skeletal code S.IV, square 2 (F2320)) and carried a Y-chromosome haplogroup R1b (R-L754), the same haplogroup as the later Villabruna individual from Italy. We obtained high-quality data (1.46-fold average coverage at targeted positions) for this individual, and we estimate a low contamination rate (∼1%) based on the X-chromosomal heterozygosity rate, as males with one X-chromosome are not expected to exhibit X-chromosomal variation.

There are two additional individuals from Climente II, one of which dates to more than 1000 years after the oldest sample, but all three are Epipaleolithic, dating to before 11700 years BP. One is a male neonate (genetic ID I33593, archaeological code S.III, 0.20, 462) dated to 14004–13512 years cal BP (12345±44 BP; after the correction for the freshwater reservoir effect: 11863±76, OxA-42701). These data (0.25-fold average coverage) do not appear to be highly contaminated (estimated at 3±0.4% by AuthentiCT^51^). The Y-haplogroup is H-P96 which has been found in later Neolithic populations, especially in the Balkans, and persists at low levels across Western Eurasia today. The other is an adult male dated to 11690–11192 cal BP (9899±79 BP, after the correction for the freshwater reservoir effect: 10425±40, PSUAMS-12555). This individual (genetic ID I17060, skeletal code #1) has higher-quality data than the oldest sample, with an average coverage of >9-fold at targeted positions, and an X-chromosome-based contamination rate estimated at ∼0.5–0.8%. He shares the same R1b haplogroup as the oldest individual.

The next individual is from Franchthi Cave on the coast of the Argolid Peninsula in southern Greece. We made multiple attempts to obtain a radiocarbon date from this individual (Fr 1 in the excavation records^52^), but these were unsuccessful due to insufficient collagen preservation. However, radiocarbon dates were obtained from a nearby hearth and on two other Lower Mesolithic samples (Fr 2 and Fr 6) and two other individuals from the site indicating a date of ∼10.5 kya^53^. The individual is male and has a Y-chromosomal haplogroup of I-M436, a subclade of I2 that was common among Mesolithic hunter-gatherers and is found among modern people, especially those of European-related ancestry^21,54^. This individual has modest coverage (0.181x) and a high estimated rate of contamination of 10–15% (95% confidence interval) based on the rate of variation on chromosome X^55^. Using *qpAdm* to model the full Franchthi data as a mixture of the damage-restricted and thus likely authentic^56^ data, and present-day Greek data (representing putative contamination), we estimate a 95% confidence interval of ∼1–11% present-day Greek-related contamination. This overlaps the estimate of contamination from heterozygosity on the X-chromosome, if we allow for the possibility that the contamination was from a female, which would provide two contaminating X-chromosomes.

Also in Greece, we have two individuals from Maroulas, an open-air site on the island of Kythnos in the central Aegean. The first Maroulas individual is a female radiocarbon-dated to 8992–8650 cal BP (7970±30 BP, UGAMS-72089), with high-quality data (1.37-fold at targeted positions). As she is female, we lack a Y-haplogroup and cannot directly estimate contamination using the X-chromosome. We estimate a contamination rate of 1–2% based on variation in runs of homozygosity on the autosomes^55^, and this individual’s mitochondrial match rate to the consensus sequence is high, again indicating low contamination (95.9–98% match to consensus – 95% confidence interval). We also present a second genome from Maroulas, deriving from an individual who lived several centuries later, and directly dated to 8594–8430 cal BP (7750±30 BP, UGAMS-72090). This individual was a determined by X and Y chromosomal ratio to be male who died as a child. His data have ∼0.81-fold coverage at targeted positions, and his Y-haplogroup appears to be a basal IJ lineage upstream of I. This individual has multiple derived haplogroup-I calls as well as multiple explicitly ancestral haplogroup-I calls. This would not be expected if missingness were driving the signal, but could be caused by contamination from an individual with a non-I Y-haplogroup. This individual has substantial contamination (∼20% from X-chromosome variation).

After this, we present an individual from Badanj Cave from the Herzegovina region of Bosnia and Herzegovina. This individual is a male and is directly dated to 16571–16375 calBP (13530±49 BP, OxA-41672). The data from this individual are of rather low coverage (0.060x) and are highly contaminated, with ∼39% contamination based on AuthentiCT^51^, which uses the rate of damage at the ends of reads to estimate contamination proportions.

In mainland Italy, we also present data from an individual from the southern tip of Apulia from Grotta Romanelli. She was a female and her remains date to roughly 13276–12816 cal BP (11110±110 BP, OxA-X-3144-12). Her genome is of moderate quality with 0.218-fold coverage at targeted sites, and a low level of contamination (∼2.5% inferred by AuthentiCT^51^).

Finally, we present Mesolithic data from the Sardinian site of S’Omu e S’Orku: a male AMS-dated to roughly 9446–9080 cal BP (8226±27 BP OxA-42320). His Y-haplogroup is I-S2555, found across Mesolithic Europe, and still present in Europe today. His genomic coverage is moderate (∼0.167-fold at targeted positions), and as such, we can use his X-chromosome to estimate that the individual is only marginally contaminated, with roughly 0.5–4.1% of his data attributable to contamination (95% confidence interval). Based on his mtDNA, which has a U5b1 consensus haplotype, the match rate to the consensus sequence is estimated at 94.6% to 98.3%, again suggesting a low contamination rate.

We performed analyses on a set of SNPs selected to minimize biases across different ancient DNA data generation methods (1240K, Twist, and shotgun)^57^. Data generation and extraction followed standard protocols^39^.

#### Bioinformatic processing

The bioinformatic processing of the DNA libraries presented in this study use the same alignment, haplotype calling, and pseudohaploid SNP calling ^58–64^, as in ref. ^39^.

We used pmdtools^56^ to detect excess C->T and G->A errors at the ends of reads to determine whether a DNA strand is from a genuinely ancient molecule or a modern contaminant. We then restrict to molecules with |Z| > 2.7 confidence of being genuinely ancient according to this software. Due to the fact that this reduces power (as many real molecules are also lost by this process), we tend to analyze both the full and “damage restricted” data. For datasets that do not show signals of contamination, we only use the “full data” as there is no clear need to use damage restriction, and this maximizes data.

#### PCA

We projected individuals in Extended Data 5 in smartpca^65^ using the parameters newshrink: YES and lsqproject: YES, onto a PCA space with axes formed by the modern populations, plotted on the graph.

#### Model competition with qpAdm/qpWave

To explore the history of pre-LGM and post-LGM hunter-gatherers of Europe, we use the soft-ware *qpAdm*^43^, which implements a methodology for generating unbiased estimates of ancestry proportions from populations that are clades without mixture with a proposed set of source populations (the “left populations” along with the modelled target population), which are related differentially to a set of outgroups (“right populations”). *qpAdm* does this by computing *f_4_*-statistics between all possible pairs of left populations and all possible pairs of right populations, testing whether *f_4_*-statistics including the target population are consistent statistically with being mixtures of *f_4_-*statistics involving the source populations, and interpreting the mixture coefficients as ancestry estimates, whose standard errors we can compute in a valid way with a block jackknife.

When modeling pre-Mesolithic populations, we used the following populations as right outgroups in *qpAdm* to distinguish different types of ancestries: modern Yoruba individuals from West Africa, the ∼35 kya Goyet Q116-1 individual probably from the Aurignacian culture in Belgium, ∼24 kya Gravettians from France, a ∼34 kya Upper Paleolithic individual from Muieri Cave in Romania, the ∼24 kya Siberian Paleolithic Mal’ta boy, and the ∼34 kya Upper Paleolith-ic Russian Sunghir individuals. All seven possible nonempty sets of populations were used to model the target populations, and the unused populations were added to the outgroups to provide greater leverage to distinguish ancestries and to test rigorously whether we could exclude contributions from ancestries similar to these groups. Results for all models are in the supplementary materials, and we present models using all three sources (Epi-Balkan, Spanish pre-LGM, Italian pre-LGM) in Table S3, with the simplest passing model in Figure 1 The simplest passing model is the one without negative admixture proportions, with the most populations moved to the right. If two models have positive admixture proportions and equal numbers of sources, then p-value ranking gives a relatively arbitrary winner.

When modeling Mesolithic Europeans and Central Asians, we again used *qpAdm* using a similar setup, albeit with a seven-source model. These sources are Epi-Spanish (represented by a Belgian Magdalenian from Fournol related to it), Epi-Italian (represented by Italian Epigravettian individuals), Epi-Balkan (represented by Romanian Epigravettians from Climente II), Anatolian LGM ancestry (represented by the Pınarbaşı individual), Caucasus LGM ancestry (represented by individuals from Kotias in Georgia), Paleolithic Siberian (represented by the Mal’ta individual, and East Asians (represented by South Chinese Neolithic islanders). To distinguish these populations, we used the following set of reference populations: modern Yoruba as a “Base” right population which should be a true outgroup to all other sources and reference populations, the Aurignacian from Goyet which shows continuity with later Magdalenian populations (Goyet Q116-1) in order to help distinguish the Spanish refugial ancestry, the Paleolithic individual from Dzudzuana Cave in Georgia, a late Paleolithic individual from Satsurblia in Georgia, a slightly older Italian Epigravettian from Sicily, whose southern location should make it less likely to be the ancestor to northern populations, pre-LGM Gravettians from Italy, Hoabhinhians from Laos, pre-LGM Aurignacians from Romania, and the Siberian Yana individuals from Siberia^66^. We also moved sources into the right reference panel if they were not required in the model. For each target population, we tried every combination of 1, 2, 3, 4, 5, 6, and 7 source models, yielding 127 non-empty sets of sources. We moved unused sources to the right to make model rejection more robust. Results for the “best” model, that is, the passing model without negative admixture proportions with the most populations moved into the right (and p-value ranking determining ties), are shown in Figure 1, and Extended Data 3, and results for every possible model are in Tables S5-S8. Populations with no fitting models are not plotted, but can be found in the supplement, and mostly consist of populations that require >100% ancestry from some source. For example, the Sardinian Mesolithic require >100% Epi-Italian ancestry, which makes sense given their higher Italian pre-LGM ancestry as seen above in Figure 1 and Tables S5-S9). For faster computation, we ran qpWave/qpAdm on precomputed output from qpfstats runs (https://github.com/DReichLab/AdmixTools/blob/master/qpfs.pdf) with a poplistname that includes Han.DG, and all target, source and right populations, and parameters allsnps: YES, inbreed: NO.

#### Y-chromosome haplogroup inference

We used the methodology described in ref.^67^, which employed the YFull YTree v.8.09 phylogeny (https://github.com/YFullTeam/YTree/blob/master/ytree/tree_8.09.0.json), and denote Y-chromosome haplogroups in terminal notation^68^.

#### Estimates of dates of admixture

We used DATES^38,44^ to estimate admixture dates for the two Maroulas individuals. We used various sets of European Mesolithic populations and Anatolian Epipaleolithic and Neolithic populations as the sources. It is more important to use many source samples even if they are somewhat genetically drifted from the true ones; picking the wrong sources does not bias the date estimate^44^.

#### Radiocarbon dating

As part of this study, samples were submitted for radiocarbon dating to three accelerator mass spectrometry (AMS) laboratories.

Measurements OxA-40804 and OxA-42701 from Climente II (Romania), OxA-41672 from Badanj (Bosnia and Herzegovina), OxA-42320 from S’Omu e S’Orku (Italy), and OxA-X-3144-12 from Grotta Romanelli (Italy) were processed at the Oxford Radiocarbon Accelerator Unit (ORAU), University of Oxford. Collagen was extracted following Law and Hedges^69^, followed by the revised gelatinization and filtration protocol of Bronk Ramsey et al.^70^, and dated by AMS as described by Bronk Ramsey et al.^71^.

AMS measurement PSUAMS-12555 from Climente II (Romania) was processed at the Penn State Radiocarbon Laboratory (PSUAMS), The Pennsylvania State University. Bone collagen was extracted and purified using a modified Longin method^72^, with gelatinization followed by ultrafiltration of the >30 kDa fraction^73^, and dated by AMS.

Two bone bioapatite AMS measurements, UGAMS-72089 and UGAMS-72090, from Maroulas (Greece) were processed at the Center for Applied Isotope Studies (CAIS), University of Georgia. Bioapatite was pretreated with acetic acid to remove secondary carbonates following Cherkinsky^74^. The purified carbonate fraction was acidified to produce CO, graphitized and dated by AMS.

Diet-derived reservoir offsets affect radiocarbon measurements on Mesolithic and Neolithic human skeletal remains from the Danube Gorges/Iron Gates region^75,76^ and require correction of the measured radiocarbon ages. For the AMS measurements obtained on the Climente II individuals, diet-derived freshwater reservoir offsets were estimated from δ¹ N and δ¹³C values using the approach outlined by Cook et al.^75^.

Radiocarbon ages were calibrated using the IntCal20 calibration curve^77^ in OxCal v.4.4^78–81^. Calibrated ranges cited in the text are reported following the conventions originally recommended by Stuiver and Polach¹³, adapted to the increased precision of more recent datasets. Range endpoints were rounded outwards to the nearest 10 years because the measurement uncertainties exceed 25 radiocarbon years.

#### Stable isotope analysis

Stable isotope dietary analysis comprised measurements of carbon (δ¹³C) and nitrogen (δ¹ N) isotope ratios. Collagen was extracted and analysed at the Oxford Radiocarbon Accelerator Unit (ORAU), University of Oxford, following the procedure outlined by Privat et al.^82^. Carbon and nitrogen stable isotope measurements were obtained using a SERCON 20-22 isotope-ratio mass spectrometer (IRMS) coupled to a SERCON GLS combustion unit.

Samples were analysed using three-point calibration with alanine, cow collagen and seal collagen standards. These in-house standards are regularly calibrated against the international USGS40 and USGS41 glutamic acid reference materials^83^. Carbon and nitrogen isotope values are reported in delta notation (‰) relative to the internationally accepted VPDB and AIR scales, respectively. Analytical precision was better than ±0.2‰ for both δ¹³C and δ¹ N.

Additional details, per sample, on C14 dating and isotopic analysis can be found in Table S11.

#### Ethics statement

All individuals from whom novel data were obtained were analyzed using methods designed to minimize damage to their skeletal remains, with authorization obtained from the relevant local authorities in each place of origin. To uphold the principles of open science, we make publicly available not only the digital molecular records (the uploaded sequences), but also the molecular records themselves (the ancient DNA libraries, which serve as repositories of molecular information). Researchers interested in conducting deeper sequencing of the libraries reported in this study may submit a request to the corresponding author D.R. Reasonable requests will be accommodated for as long as the libraries remain preserved in our laboratories, and such access will not be contingent on including us as collaborators or co-authors on any resulting publications.

#### Reporting summary

Further information on research design is available in the Reporting Summary linked to this article.

## Data availability

Genotype data for individuals included in this study are available from the Harvard Dataverse repository at https://doi.org/10.7910/XXX/XXXXXX. The DNA sequences reported in this paper have been deposited in the European Nucleotide Archive under accession number PRJEBXXXXXXXX. Other newly reported data, such as radiocarbon dates and archaeological context information, are included in this paper and the Supplementary Information.

## Code availability

No novel code is needed to replicate the results of this study. The ADMIXTOOLS package can be downloaded from https://github.com/DReichLab/AdmixTools.

## Acknowledgments

We acknowledge the ancient and modern individuals whose data we used for analysis. We thank the museum curators, laboratory technicians, and other individuals who may have directly or indirectly helped generate the data we used. We acknowledge the system administrators and funders who help maintain the Harvard Medical School O2 research computing platform, where these analyses were performed. We thank Nicole Adamski, Rebecca Bernardos, Kim Callan, Jeremy Choin, Trudi Frost, Ilana Greenslade, Aisling Kearns, Jack Kellogg, Ann Marie Lawson, Matthew Mah, Adam Micco, Mariam Nawaz, Noah Workman, Lijun Qiu, and Fatma Zalzala for help with sample preparation and or critical comments on the manuscript.

## Author Contributions

D.T., D.B., A.P.S., C.B., M.M., R.M., N.R., I.L., C.Posth, M.H., E.B., and D.R. wrote the manuscript and created the supplementary information; D.T. and A.P.S. analyzed the data; D.T. Interpreted the results; D.T. generated all figures with input from A.P.S., C.B., E.B., and D.R.; D.B., I.L., N.R., S.M., N.P., C.Posth, M.H., E.B., and D.R. supervised various aspects of the work; D.B. compiled the archeological aspects of the work; D.B., C.B., T.C., F.A, A.B., E.C., F.C., C.M.F., A.L., A.Marić, A.Masciana, M.M., R.M., A.P., C. Perlès, R.P., T.D.P., K.K., A.Sampson, A.Soficaru, S.K., A.T., K.U., and O.E.Y. excavated, curated and or shared skeletal materials and the archeological information; D.T., A.P.S., G.S., N.R., S.M., and D.R. performed bioinformatic data processing and curation and contamination estimation; A.P.S., N.R., and M.H. carried out or supervised wet laboratory work; D.T. and D.B. conceived of the study.

## Additional Information

Supplementary Information is available for this paper.

Correspondence and requests for materials should be addressed to D.T., D.B., or D.R.

---

**Extended Data 1:**
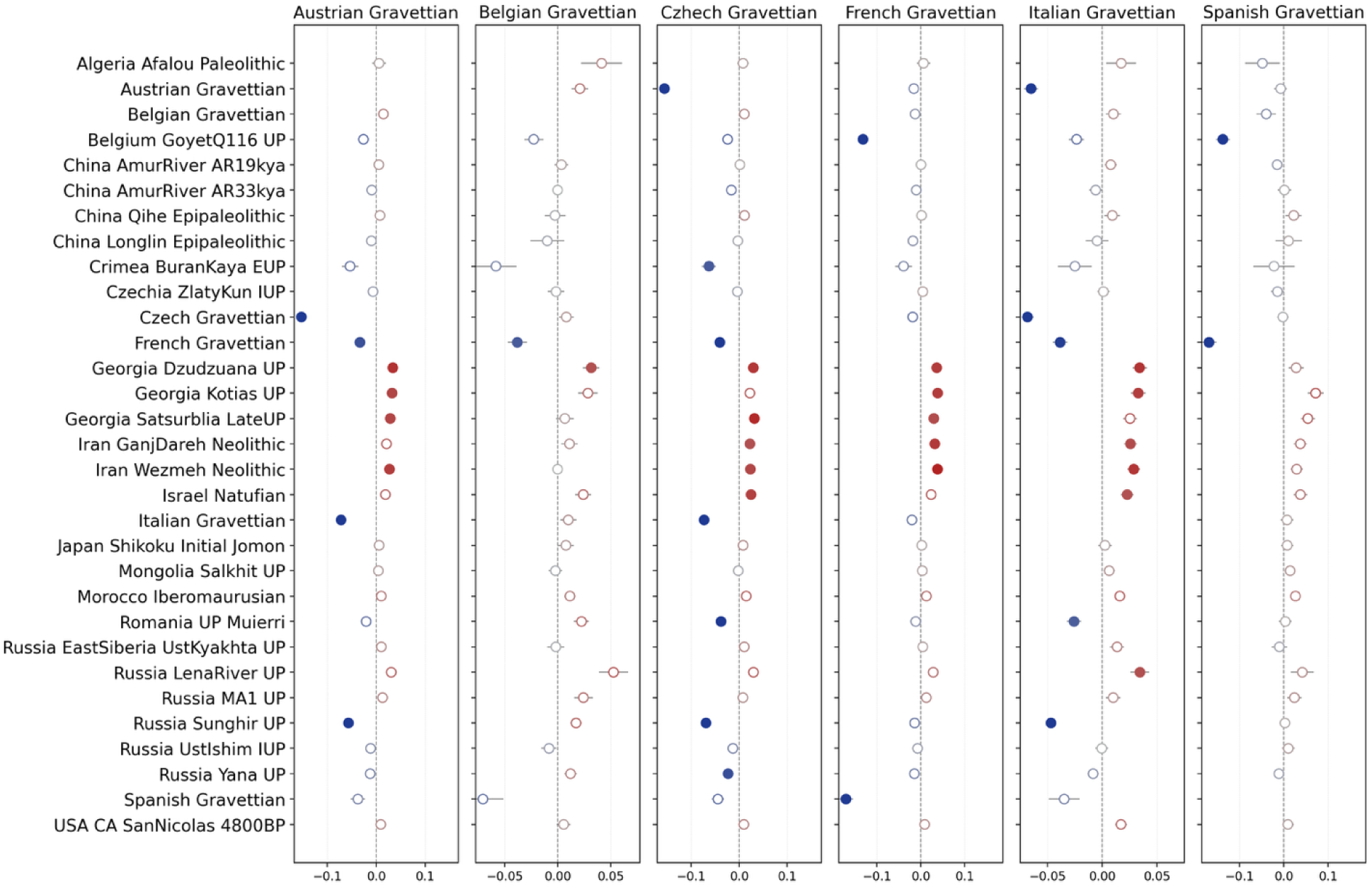
No evidence for excess Siberian or East Asian ancestry in Epi-Balkan people, but strong evidence for Near Eastern affinity. Z-scores for *f_4_*-statistics of the form *f_4_*(Outgroup, Row population; Column population, Epi-Balkan), where the row populations are a diverse set of Eurasians which are generally older than Epi-Balkan, and where the column populations are a diverse set of Pre-LGM Gravettians from across Europe. Statistics with |Z|>4 are red for positive *f_4_*s, or blue for negative. Standard errors are plotted. In agreement with previous studies^5,13,49^ based on other post-LGM, pre-Neolithic Europeans, the Epi-Balkan population has an affinity to Near Eastern groups, while lacking any evidence of Basal Eurasian ancestry. In tension with previous studies^5^ the Epi-Balkan population shows no excess affinity to the oldest East Asian and Paleolithic Siberian populations, potentially suggesting that movement of Epi-Balkan-related ancestry eastward could explain the affinity seen in later East Asian, Siberian, and Native American groups.

**Extended Data 2:**
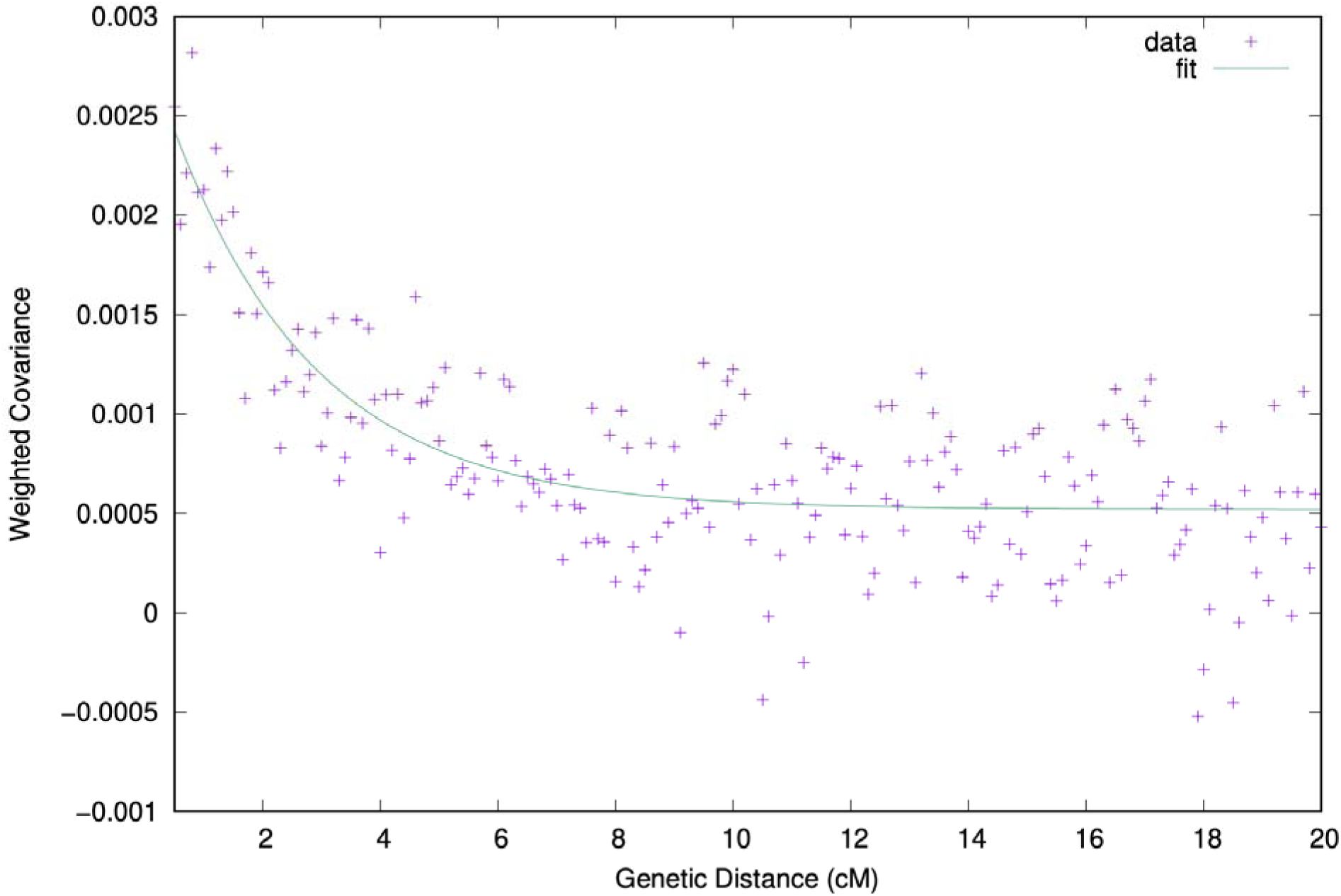
DATES output. DATES curve for the pool of two Maroulas individuals, modeled as mixtures of the Serbia Iron Gates Mesolithic and Anatolia Barcın Neolithic as admixture sources. The decay implies mixture 1034±266 years before the dates of the Maroulas individuals (Z > 3.7 for evidence of a decay).

**Extended Data 3:**
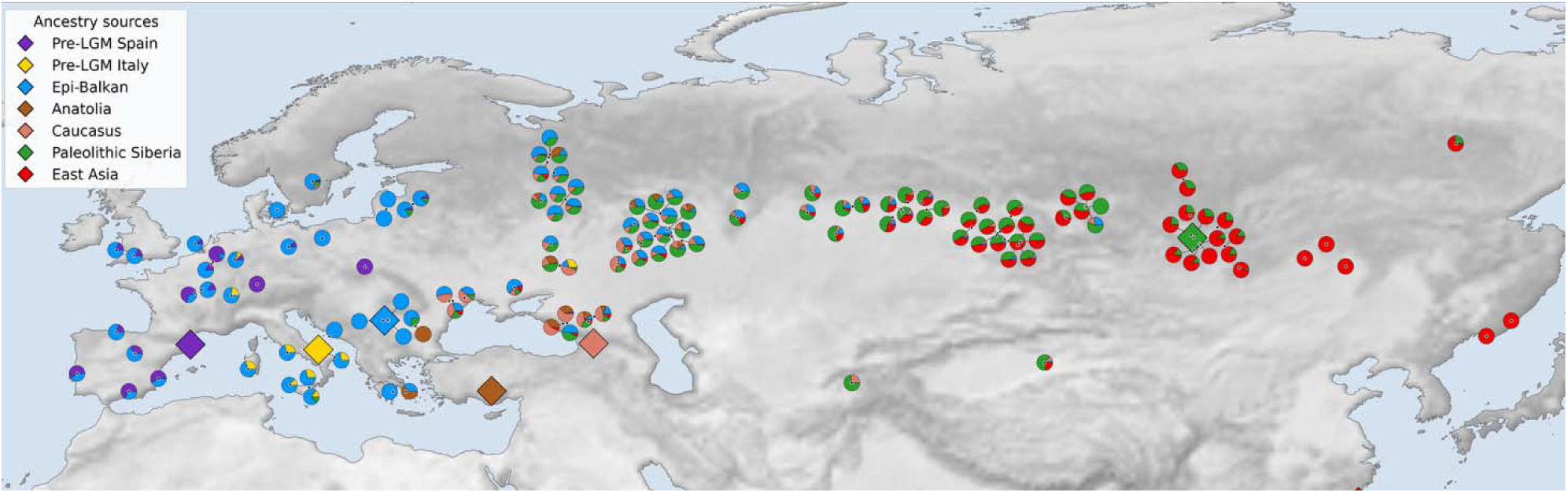
Seven Source Model of Mesolithic Eurasia. Modeling much of Eurasia with seven ancestry sources found across Europe, Eurasia, Central Asia, and Siberia. The simplest, passing, non-negative model with sources rotated into the right is the one plotted. The large diamonds represent the locations of source populations.

**Extended Data 4:**
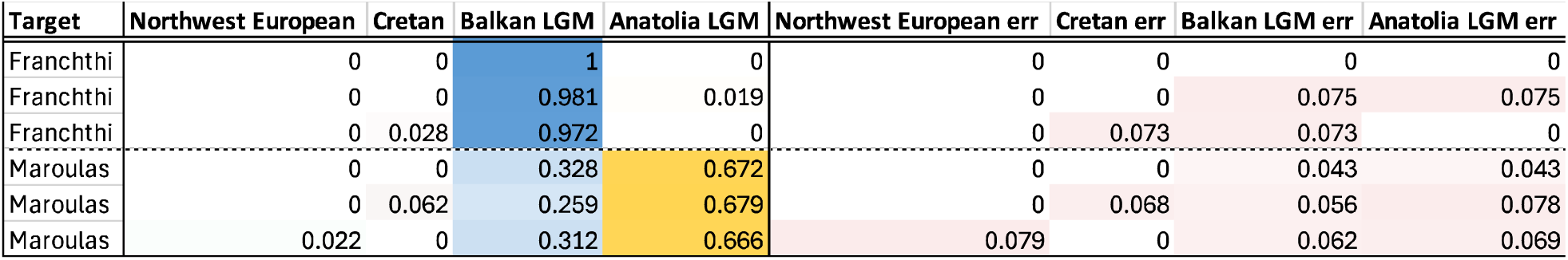
Maroulas Individuals are of mixed Balkan and Anatolian ancestry. The Maroulas individuals can only be modelled as a mixture of Epi-Balkan and Anatolian Epipaleolithic ancestry, even when Northwestern Europeans (represented by European Americans from Utah - CEU) and modern Cretans are provided as possible sources. Franchthi can only be modelled as being a clade within the Epi-Balkan population, in line with previous findings. Only models which have p > 0.01 and non-negative admixture proportions are displayed. All unused left populations are rotated into the right, joining the following set of base populations: Yoruba, Goyet-Q116-1, Dzudzuana, Kotias, Natufian, Yamnaya, and Sunghir.

**Extended Data 5:**
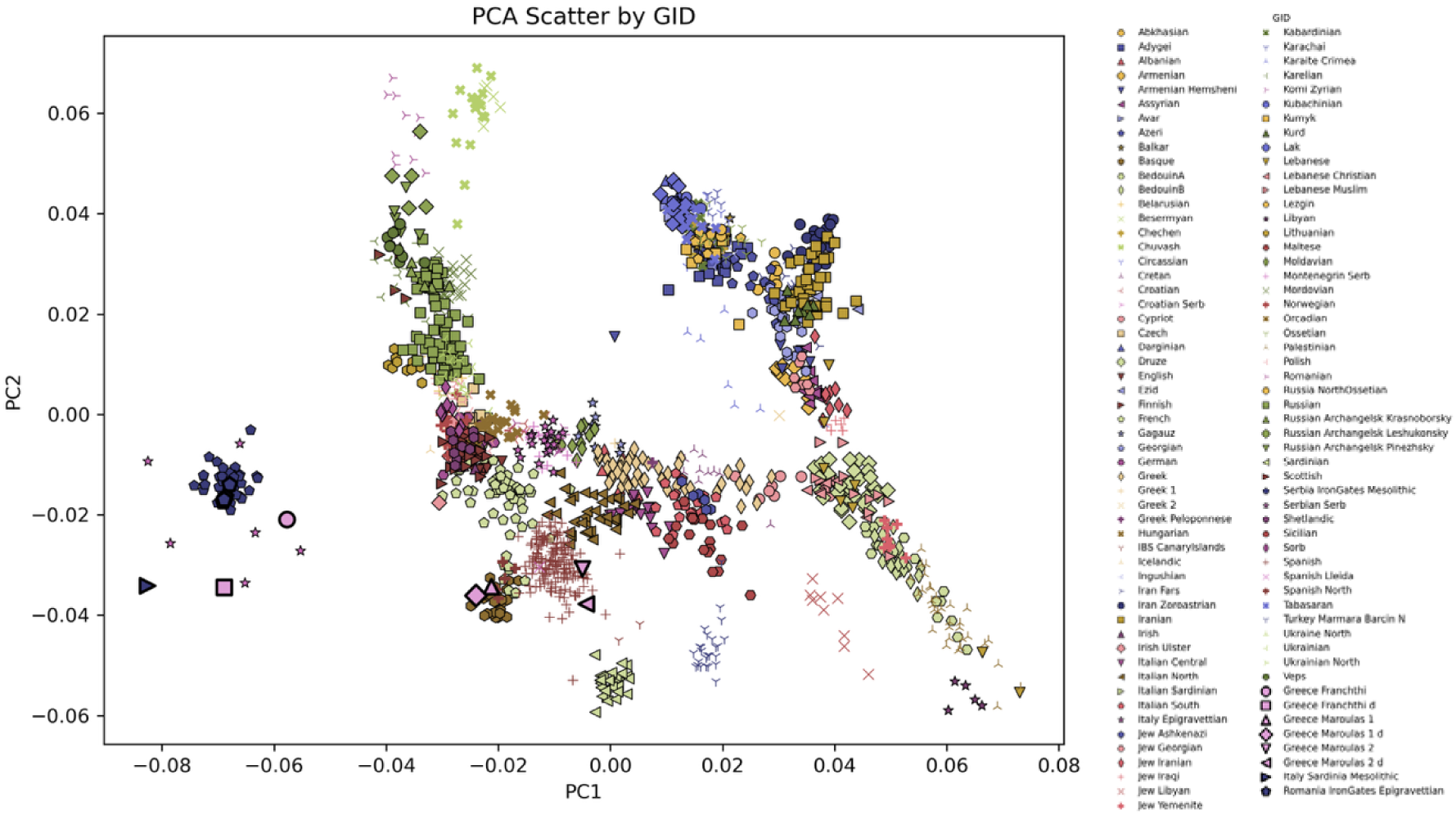
West Eurasian PCA shows that Mesolithic Greek samples lie between Mesolithic Hunter-Gatherers and Neolithic farmers. The standard West Eurasian PCA shows that Greek Mesolithic samples (represented by the large pink square and circle (Franchthi), as well as diamond and triangles (Maroulas individuals) appear to be positioned between Mesolithic European and Neolithic populations. This reflects gene flow between Anatolia and Mesolithic Europe.

