## Supplement for "The Pan-European Impact of the Balkan Hunter-Gatherers"

**Supplementary text – archaeological sites and context descriptions**

**Climente II (Danube Gorges, Romania)**

Climente II Cave (44° 38′ 10″ N, 22° 17′ 3.99″ E) was situated on the left (Romanian) bank of the Danube in the Lower (Kazan) Gorge of the Danube Gorges region, approximately 12 m above the original river level (Fig. S1–3). The site was excavated by Vasile Boroneanț in 1968–1969. An Epipaleolithic occupation was identified within the cave, consisting of a quasi-circular hearth, numerous lithic artifacts, bone and antler tools and weapons, personal adornment beads, and faunal remains.

The site also yielded the remains of an articulated primary inhumation lacking the cranial bones and one of the scapulae. The individual had been placed in a flexed position on the left side (Fig. S4). The lower limbs were tightly flexed, with the feet positioned behind the pelvic area. The forearms were bent at the elbows, and the hands were placed in front of the face. The body had been placed in a natural depression in the bedrock. Ochre was observed in the area where the skull had been, and on and below the postcranial bones. Two direct AMS radiocarbon dates obtained for this individual and reported by Bonsall et al. (2012, 2016) place the burial within the Bølling–Allerød interstadial (combined OxA-22042 and OxA-24990: 12220 ± 58 BP), calibrating to 14790–13888 cal BP at 95% confidence after correction for the Danube reservoir effect. A few bone fragments and several teeth were found in Trench S III, a few meters nearer the cave entrance in a shallow water-worn gully. The excavator assumed that flooding had washed away the missing parts of an originally intact skeleton.

The genetic data reported in this article derive from three individuals. The first individual is represented by a molar from an adult and comes from Trench S IV, square 2 (F2320). The molar is directly dated to 12687 ± 45 BP (OxA-40804). The associated isotopic values, δ^13^C = –18.2‰ and δ^15^N = 13.7‰, indicate a freshwater reservoir effect that requires an offset correction (Cook et al. 2002). After correction, the date is 12349 ± 63 BP, which calibrates to 14842–14100 cal BP. Microbiome genetic data (Ottoni et al. 2021) and strontium isotope data (Borić and Price 2013) have been obtained from the same tooth. The second individual is represented by a neonate tibia covered in red ochre (Fig. S5), with stratigraphic details recorded as S. III, 0.30, 962. It was directly dated to 12345 ± 44 BP (OxA-42701) and is associated with isotopic values of δ^13^C = –18.5‰ and δ^15^N = 18.6‰. The highly elevated δ^15^N value likely indicates a nursing signal, and this measurement requires application of the maximum correction factor of –545 ± 70 years. (In human bone collagen studies, exclusively breastfed infants typically show a δ^15^N offset of approximately 2.0‰ to 3.0‰ higher than their mothers. Accordingly, we subtracted 2.5‰ from the neonate δ^15^N value (= 16‰) before applying a reservoir correction.) After correction, the date is 11863 ± 76 BP, which calibrates to 14004–13512 cal BP at 95% confidence. The third individual is represented by a second molar (M2) and is directly AMS-dated to 10425 ± 40 BP (PSUAMS-12555). The reservoir-corrected age, based on the δ^15^N value of 16.7‰, is 9899 ± 79 BP, yielding a calibrated age range of 11690–11192 cal BP at 95% confidence during the Early Holocene.

*References*: Bonsall et al. 2012; 2016; Borić and Price 2013; Boroneanţ, A. 2011; Boroneanţ, V. 1970, 2000; Cook et al. 2002; Ottoni et al. 2021.

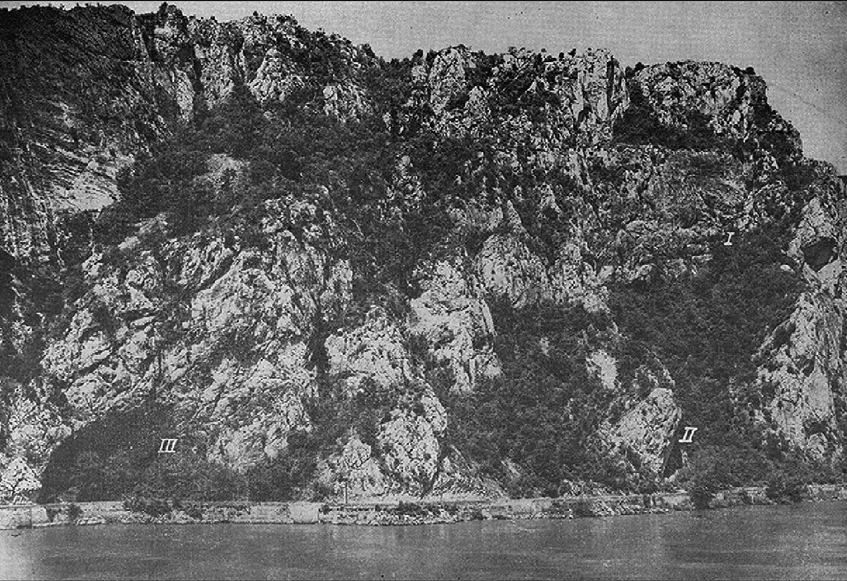

Fig. S1. Caves opening in the Ciucaru Mare escarpment (I – Climente I, II – Climente II, III – Cuina Turcului) (photo: Institute of Archaeology in Bucharest).

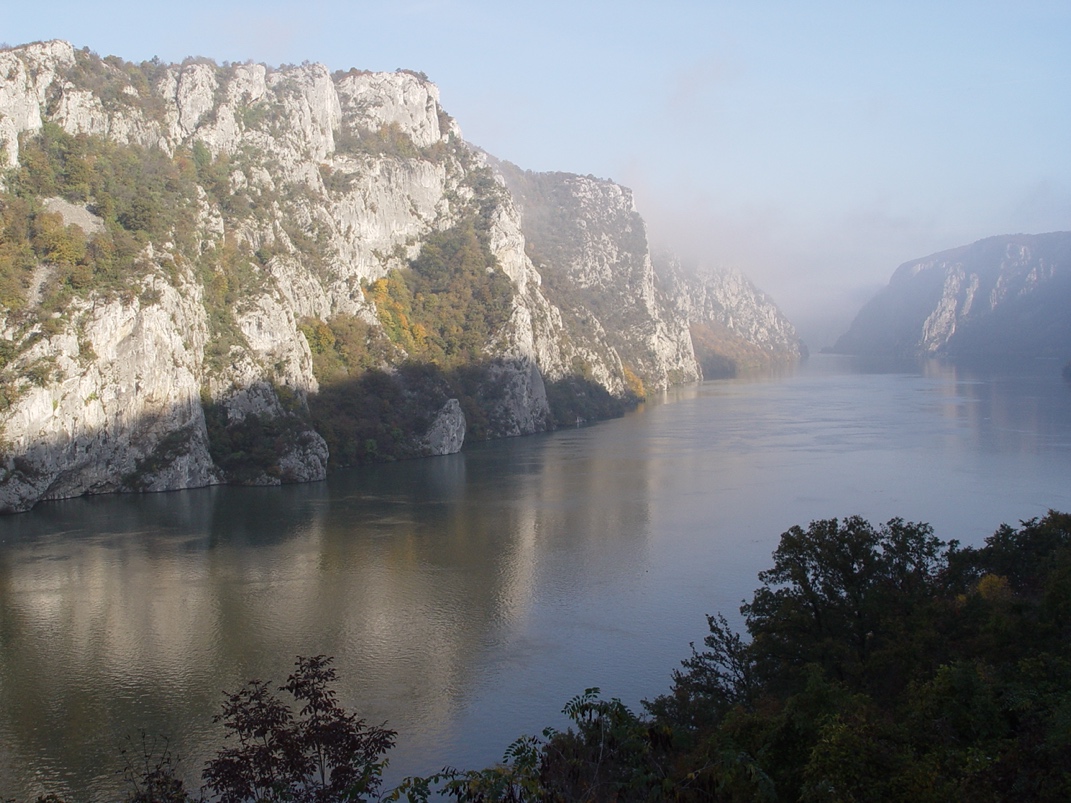

Fig. S2. Ciucaru Mare escarpment today within the Kazan Gorge of the Danube Gorges area (photo. D. Borić).

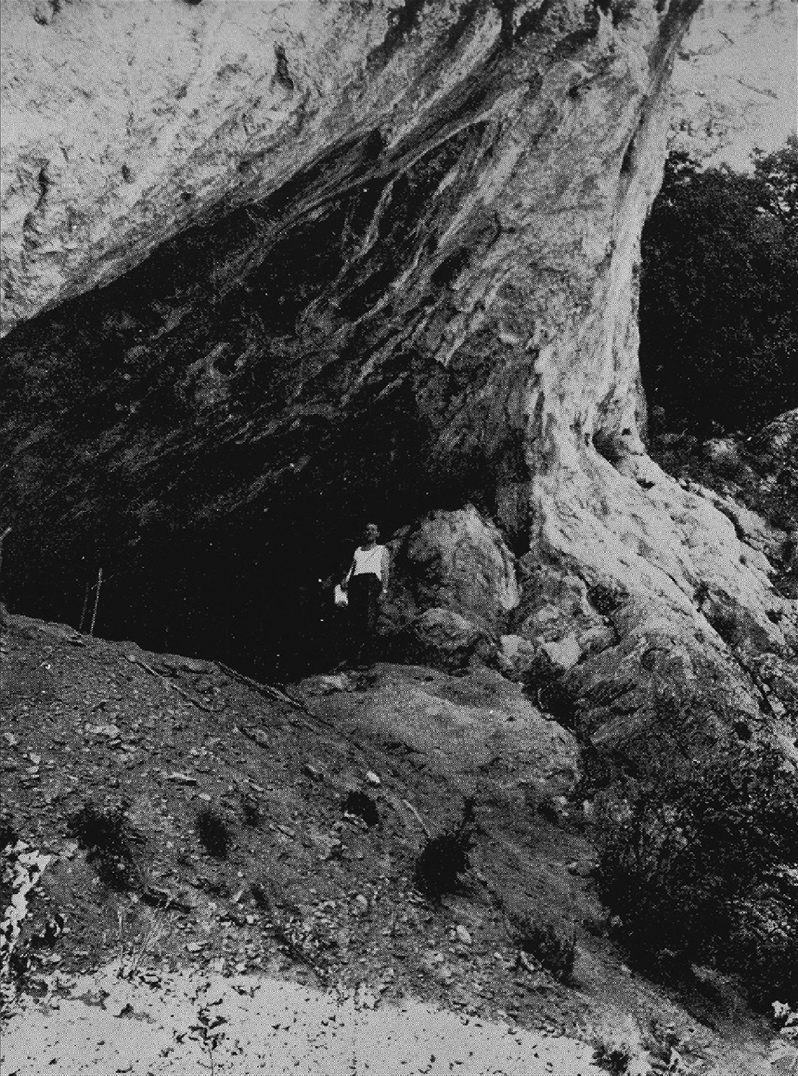

Fig. S3. Entrance to Climente II Cave (photo: Institute of Archaeology in Bucharest).

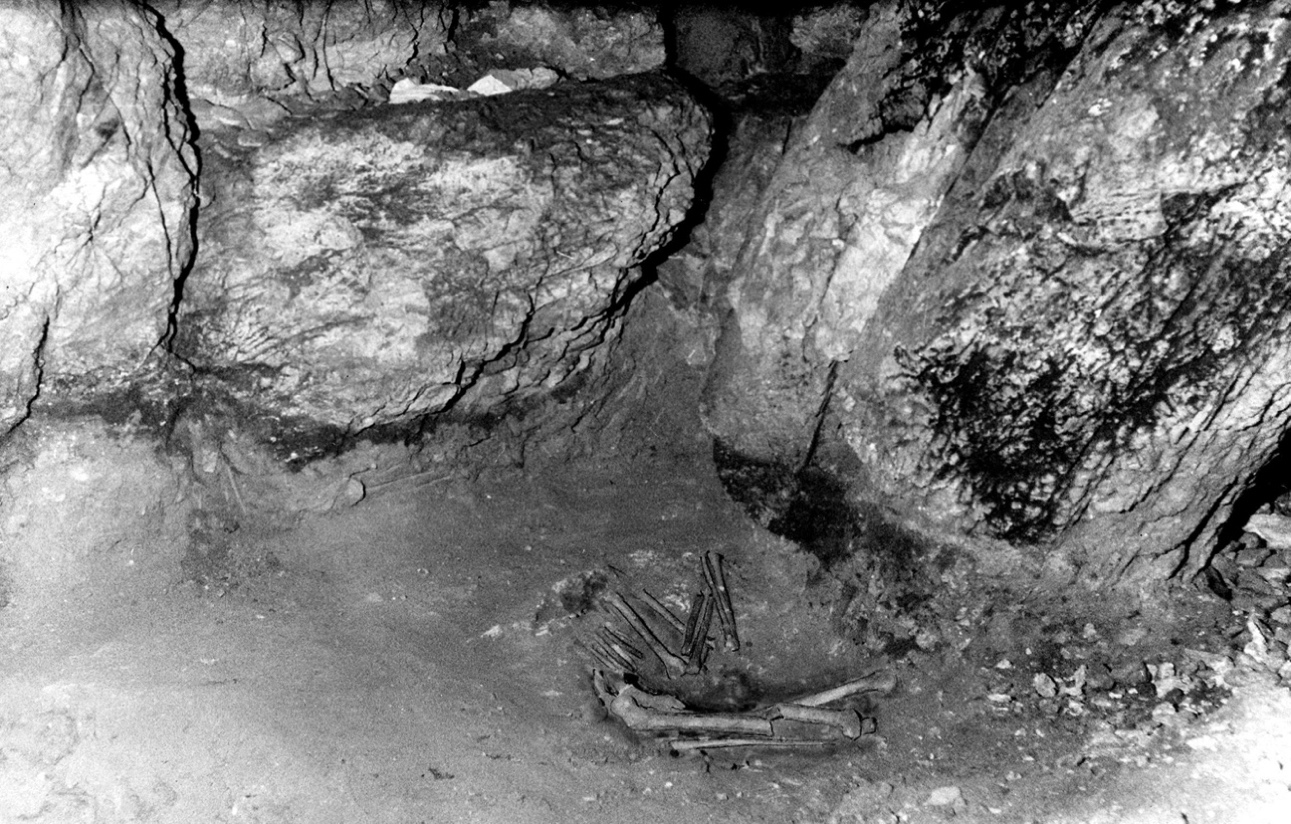

Fig. S4. Climente II burial M1 *in situ* (photo: Institute of Archaeology in Bucharest).

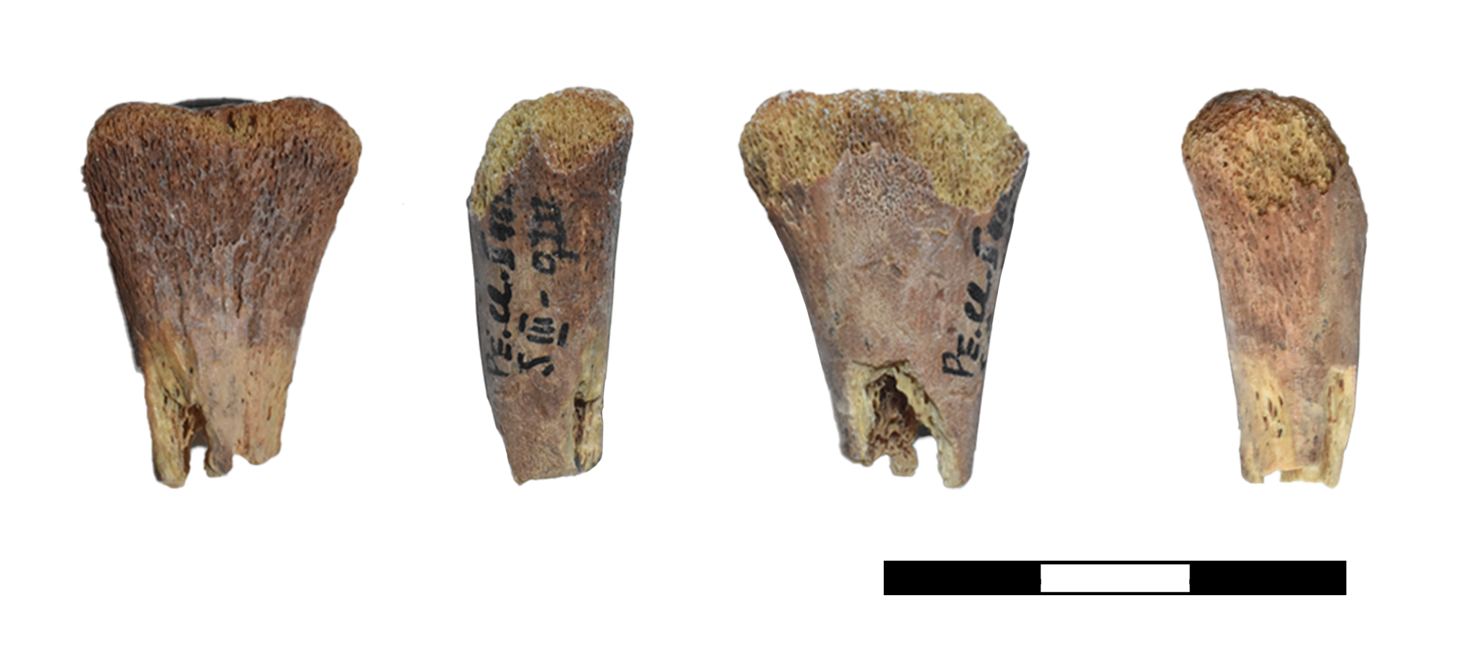

Fig. S5. Sampled neonate bone from Climente II.

**Badanj (Herzegovina, Bosnia and Herzegovina)**

The rockshelter of Badanj (43° 4′ 52.77″ N, 17° 53′ 7.94″ E; c. 100 m asl) is located near the village of Borojevići, 7 km west of the town of Stolac, along the Bregava River. The area of the site protected from the elements covered approximately 40 m² (Fig. S6). The site is particularly well known for a rock engraving carved on a large boulder situated in the center of the eastern part of the shelter. Excavations were first conducted by Đuro Basler, curator at the National Museum of Bosnia and Herzegovina in Sarajevo, between 1976 and 1979, covering around 50 m² of the eastern part of the shelter (Basler 1976, 1979). In 1986–1987, an international team led by Zilka Kujundžić-Vejzagić and Robert Whallon investigated the western part of the shelter, exposing an area of approximately 35 m².

Based on the available radiocarbon dates, the site was occupied for several millennia during the Upper Palaeolithic, beginning during the Last Glacial Maximum around 20000 years ago and continuing into the Oldest Dryas. These early Epigravettian phases extended into the Late Epigravettian during the Bølling–Allerød interstadial, from c. 16000 cal BP to the onset of the Younger Dryas around 13000 cal BP (Borić et al. 2023; forthcoming). During the 1976–1979 excavations, the depth reached in different parts of the trench surrounding the engraved boulder ranged between 1.5 m and 2.3 m below the surface encountered in 1976. During the second phase of research in 1986–1987, approximately 29 archaeological layers were identified within a trench about 1 m deep, although in many areas the depth varied between only 80 and 90 cm.

The chipped stone assemblage is characterized in the earlier phases by a predominance of backed bladelets with straight or slightly curved backs, whereas the later phases, corresponding to the Bølling–Allerød interstadial, are dominated by thumbnail or “circular” scrapers. Backed bladelets with strongly curved backs are more characteristic of the later phases, while backed blades and both small and large backed flake points also become more frequent during this period. Geometric microliths, mainly crescents/segments with approximately half as many triangles, are restricted exclusively to the later occupation phases. End-scrapers on blades and large, robust side-scrapers decline in frequency from the earlier to the later periods, whereas truncations become more common in the later phases.

Red deer was the predominant hunted species throughout the sequence, particularly during the later and warmer phases. Ibex and chamois are well represented in the faunal assemblages of the earlier, colder phases, whereas wild boar and roe deer increase in frequency during the later occupations. Numerous lithic and osseous artifacts were recovered from the site, including engraved bone artifacts, together with ornamental beads manufactured from Tritia gibbosula, Tritia neritea, Columbella rustica, Dentalium sp., Glycymeris sp., and red deer canines (Borić et al. 2023; Whallon 2025).

During the analysis of faunal remains excavated in the 1976–1979 campaigns, a human first metatarsal (quadrant XIX/9, depth 1.3–1.6 m, recovered 26/10/1979; ZooMS 243) (Fig. S7) and a human phalanx (quadrant XIII/8, depth 1.2–1.3 m, recovered in 1977; ZooMS B262) were identified. Although attempts were made to recover ancient DNA from both specimens, only the metatarsal yielded sufficient endogenous DNA for further analyses. Based on the recorded depths below the excavation surface, both specimens are likely attributable to the earlier phase of site occupation. The AMS radiocarbon date obtained for the metatarsal, 13530 ± 49 BP (OxA-41672), was accompanied by the following stable isotope values measured during indicative AMS burns: δ^13^C = –19.0‰ and δ^15^N = 11.1‰. These values are consistent with a terrestrial diet based primarily on C3 food pathways.

*References:* Basler 1976, 1979; Borić et al. 2023; forthcoming; Whallon 2025.

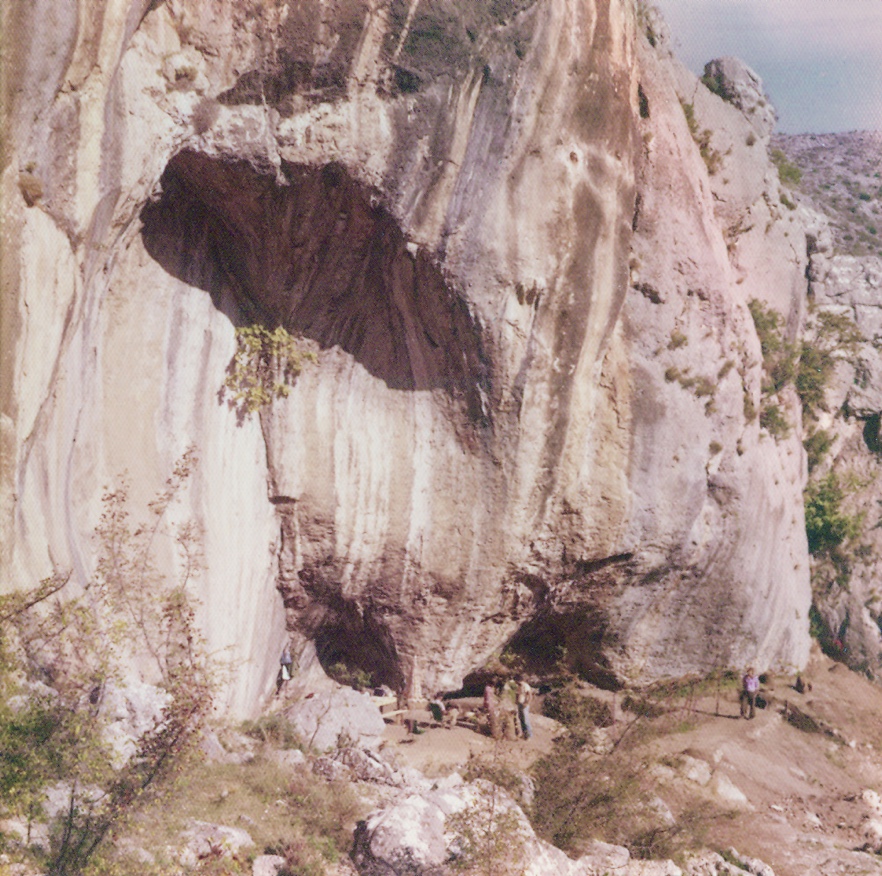

Fig. S6. Badanj rockshelter (photo: D. Šljivar, after Borić et al. 2023).

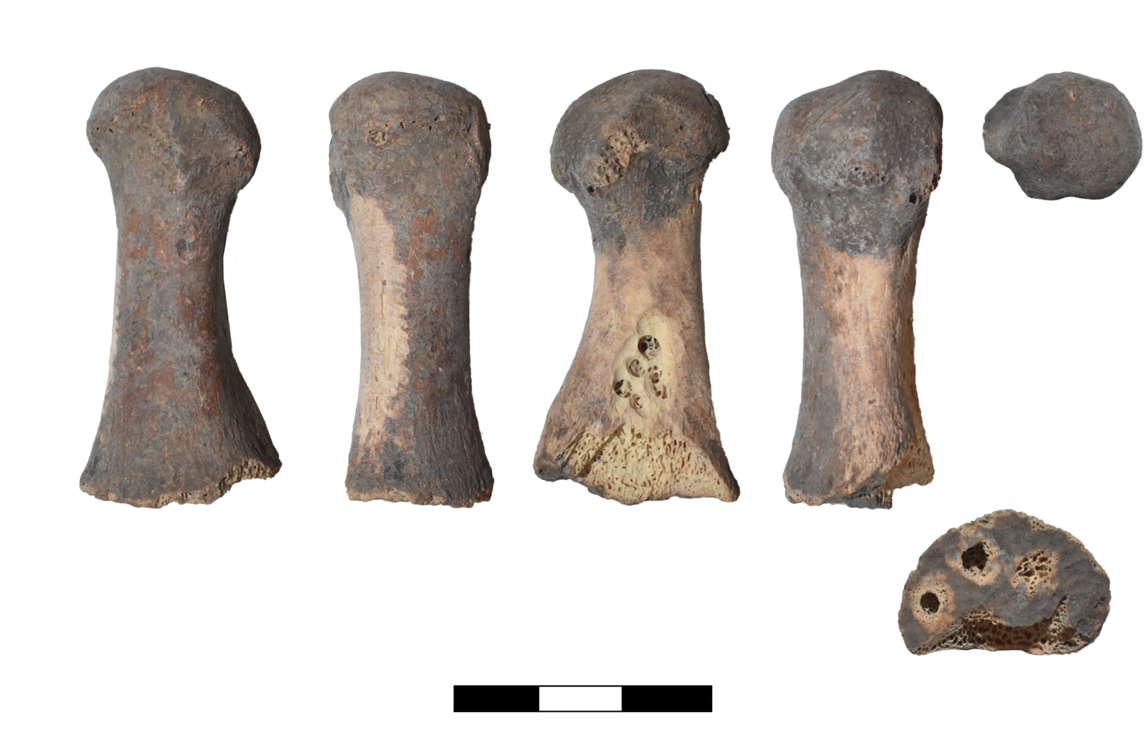

Fig. S7. Sampled metatarsal I from Badanj (photo: D. Borić).

**Franchthi (Argolid, Greece)**

Franchthi Cave (**37° 25′ 24″ N, 23° 07′ 56″ E**) is located in the Argolid directly on the present-day coast (Fig. S8). During the Upper Palaeolithic, however, the shoreline lay between 2 and 5 km away, depending upon sea level fluctuations. The cave stood at an altitude varying from approximately 50 to 130 m asl. The site was occupied as a hunting halt exploiting large game, hare, and birds throughout the Upper Palaeolithic in several discrete phases corresponding to Perlès’ Lithic Phases 0–III, or Franchthi General Phases (FGP) 0–4 (Perlès and Rose, in prep.). Following a chronological hiatus of several millennia encompassing the Last Glacial Maximum, the cave was reoccupied from around 14500 cal BP onward. This period spans the Epipalaeolithic (or Late Upper Palaeolithic) and is associated with significant changes in lithic technology compared with earlier periods, notably the appearance and relatively high frequency of microburin technique among retouched tools. This development corresponds to FGP 4-6 (Perlès 1987, Perlès and Rose forthcoming).

During this period, occupation intensity increased relative to the Upper Palaeolithic sequence, as reflected in the appearance of marine shells, large quantities of land snails, carbonized seeds, fish and hunted mammals, primarily cervids, suids, as well as fur species. FGP 4 is characterized by specific backed points produced with the microburin technique, while the first geometric microliths (isosceles triangles), produced using the microburin technique, appear in Lithic Phase V (FGP 5, from c. 13000 cal BP), According to Perlès, Lithic Phase V marks the emergence of a distinct cultural tradition, identifiable through the character of the lithic assemblage and a “broad-spectrum economy,” including abundant evidence of carbonized plant remains, that continues throughout the Mesolithic (Perlès 1999: 314). During FGP 6 (c. 12000 cal BP), Melian obsidian appears at the site for the first time, accompanied by increasing diversification in lithic tool types, including segments and trapezes, while the quantity of large mammal remains declines and the number of land snails decreases substantially.

The Lower Mesolithic occupation at Franchthi Cave corresponds to FGP 7 (Lithic Phase VII, c. 11000 cal BP) and is characterized by a dramatic shift in lithic production. Assemblages are now dominated by flakes with end-scrapers and crudely fashioned tools, alongside the near disappearance of microburin technique, geometric microliths, and backed bladelets. This phase is also characterized by massive accumulations of land snails and carbonized seeds that indicate occupations during spring, summer, and autumn.

Rare human remains (isolated bone fragments and shed milk teeth) first appear at Franchthi during the Final Palaeolithic period. During the Lower Mesolithic, an area near the present mouth of the cave in Trench G1 served as a burial ground for at least a dozen individuals (Cullen 1995; forthcoming). The remains of six disturbed inhumations and two cremations, along with scattered bones from other individuals, represent individuals from all age groups and both sexes. Two were dated to 10655–10425 cal BP (MAMS-52044: 9328 ± 28) and 10580–10375, (MAMS-52045: 9294 ± 27 BP) with the associated isotope values indicating terrestrial dietary pathways (Martinoia *et al.* 2025). Less than half a meter above this cluster lay the shallow grave of a well-preserved adult male (Fr 1), 25–30 years old. The genetic data reported in this article relate to Fr 1, who was laid out in a fully articulated contracted position with stones piled on top of the body (Fig. S9). This individual appears to have died from blows to the forehead. The grave may originally have been lined and surrounded by small stones. A radiocarbon date was obtained from a nearby ashy deposit, possibly a hearth: 11070–10175 cal BP (P-1519: 9260 ± 140 BP). No grave goods can be associated with the burial (Cullen 1995: 275–277).

FGP 8, the Upper Mesolithic, corresponds to a period when the site became a focal point for large-scale fishing expeditions targeting bluefin tuna. The sea shore was then about 2 km from the site. The lithic assemblage indicates the presence of microliths of atypical form, perhaps associated with specialized fishing gear, but these were no longer produced using the microburin technique. At the same time, the use of Melian obsidian increases. Characteristics of the lithic technology in the final Mesolithic phase, FGP 9, demonstrate similarities with the Early Mesolithic assemblages, suggesting continuity in lithic technological traditions that were repeatedly reorganized and modified in response to changing needs during the Epipalaeolithic and Mesolithic occupation of Franchthi.

FGP 10, or “Initial Neolithic” (possibly aceramic), witnesses the introduction of newly available domestic resources, including wheat, lentils, and ovicaprids, all of which appear in the cave deposits. Most of the lithic assemblage demonstrates continuity with the preceding period, although pressure-flaked bladelets and trapezes may have been imported, as there is no evidence for their local manufacture. Obsidian continued to be used during the Initial Neolithic, while a few pottery sherds associated with the Early Neolithic occur in very small quantities in the upper Initial Neolithic deposits. This Initial Neolithic occupation of the cave may have been relatively small-scale. The elements of continuity in lithics, in shellfish and in ornaments suggest a mixed group of newcomers and descendants of local hunter-gatherers (Perlès 2023; but see Munro and Stiner 2015 for a contrary argument). Asouti et al. (2018) concur that the selective introduction of cereal crops from southwestern Asia around 8700 cal BP seen at Franchthi involved interactions between local foragers and newly arrived farmers, rather than a wholesale Neolithic “package.” The cave’s botanical remains offer a long-term perspective on environmental changes. In the Late Pleistocene and Early Holocene, this unique littoral paleohabitat in southern Argolid resembled southwestern Asia, with similar plant niches/biomes colonized by junipers, annual legumes, and grasses.

The Early Neolithic occupation of the cave, and especially of the external terraces known as “Paralia,” is characterized by a range of artefacts typical of the Early Neolithic across the wider region. It probably does not derive from the Initial Neolithic settlement and represents the arrival of new groups of farmers, who again interacted with the descendants of local hunter-gatherers. Lacking grave goods or secure context, burials from this period could be dated only to a broad range of Early–Middle Neolithic (Vitelli 1993). Six inhumations of neonates and infants fall into this category, most of which were found in Paralia. Subsequently, the sequence continues into the Middle Neolithic, but the Late Neolithic marks a sharp break in settlement patterns, pottery, flaked stones tools, bone tools, and ornaments, suggesting the arrival of groups with different traditions. The reoccupation during the Final Neolithic, after a long hiatus, is closer in settlement patterns to the Early and Middle Neolithic ones. Several other burials were uncovered in the Middle and especially Final Neolithic. A bone from a Final Neolithic adolescent indicated an ancestry different from northern Anatolian and Balkan populations, and closer to southern Anatolian ones (Mathieson *et al.* 2018)*.*

*References*: Asouti et al. 2018; Cullen 1995, forthcoming; Perlès 1987; 1999; 2001; 2023; Jacobsen 1969; 1976; Munro and Stiner 2015; Vitelli 1993; Mathieson et al. 2018; Martinoia et al. 2025; Perlès and Rose, forthcoming.

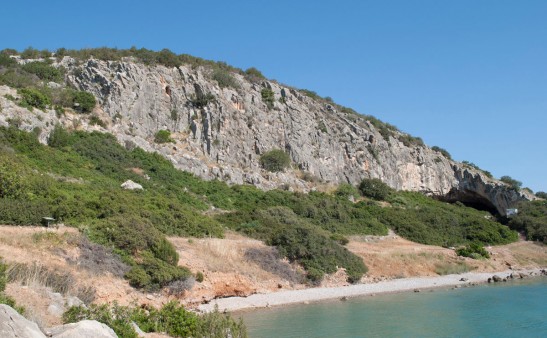

Fig. S8. Position of Franchthi Cave.

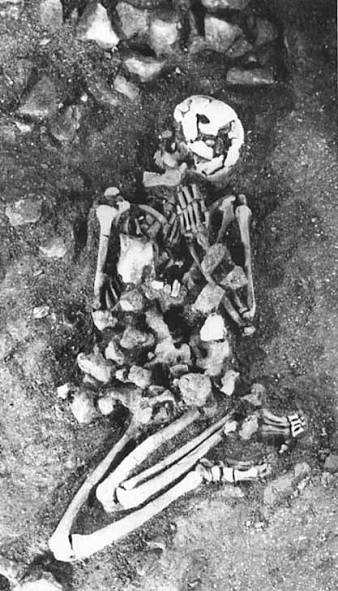

Fig. S9. Lower Mesolithic articulated Burial Fr 1 (after Cullen 1995: Fig. 3).

**Maroulas (Kythnos, Greece)**

Maroulas (37° 26′ 50″ N, 24° 25′ 53″ E) is located on an eroded promontory off the northeastern coast of the island of Kythnos, Greece, only 5–10 m above present-day sea level (Fig. S10–11). It is one of the rare open-air Mesolithic sites in the Cyclades and the wider Aegean region. The first rescue excavations at the site began in 1996 under the direction of Adamantios Sampson, while more systematic work was carried out between 2001 and 2005 through Sampson’s collaboration with Janusz Krzysztof Kozłowski (Sampson et al. 2010). The excavated area covered 2000 m² and comprised 31 trenches. More than a dozen circular and ellipsoidal limestone structures (3–4 m in diameter), interpreted as habitation dwellings, were identified and contained vertically placed stone slabs. Some structures contained articulated human burials beneath and above stone-paved floors, as well as disarticulated human remains representing at least 26 individuals within the structures’ infill deposits, often covered by stone slabs. Although most burials are single inhumations, multiple burials are also present (e.g. burials 21–23 in structure C21), together with evidence for secondary mortuary practices. Obsidian artifacts, fish vertebrae, and seashells were found around Burial 1, which had been placed in a highly contracted supine position. The burial assemblage spans a wide range of age groups, from neonates to adults.

The most up-to-date absolute chronology for the deposits is based on paired dating of marine shells and charcoal samples (five pairs in total), undertaken to estimate the effects of the marine reservoir offset on shell dates (Facorellis et al. 2010). Measurements were conducted at the Keck Carbon Cycle AMS Facility, University of California, Irvine. Two calculated means for different periods yielded ΔR values of –46 ± 73 and –238 ± 48 years. The charcoal dates obtained at the Keck Carbon Cycle AMS Facility correspond well with charcoal dates measured at the Radiocarbon Laboratory of Poznań University: square 11, spit 2: 9440 ± 40 BP (Poz-2200); trench 3, spit 4: 9420 ± 50 BP (Poz-6486). These measurements suggest occupation of the site between c. 10800 and 10500 cal BP.

Samples for aDNA analyses and AMS dating were collected under permit no. ΥΠΠΟΑ/ΓΔΑΠΚ/ΔΣΑΝΜ/ΤΕΕ/Φ77/167587/114450/1894/126, issued by the Ephorate of Palaeoanthropology-Spelaeology of the Hellenic Ministry of Culture on 2 June 2020. Collagen preservation in the samples submitted for AMS dating was extremely poor, and all nine submitted samples were withdrawn following %N pre-screening. Nevertheless, marginal but authentic endogenous aDNA was recovered from Sample #M2: Tetr. Δ1a, 5 (13/09/1996) (Fig. S12), corresponding to the right petrous portions of the temporal bone, and from Sample #M4: burial 16 (sq. 13, trench 2) (Fig. S13), corresponding to the left petrous portions of the temporal bone. Two direct bioapatite dates were subsequently obtained on these individuals at the AMS laboratory of the University of Georgia: I23419: 7970 ± 30 BP, UGAMS-72089 and I23420: 7750 ± 30 BP, UGAMS-72090, which calibrate in the approximate range of 9000–8400 cal BP. If taken at face value, these dates may suggest that some of the site’s features are younger than previously indicated by charcoal and shell dates. Consequently, further work is needed to reconcile the site’s chronology by applying additional dating proxies.

*References:* Facorellis et al. 2011; Sampson et al. 2010.

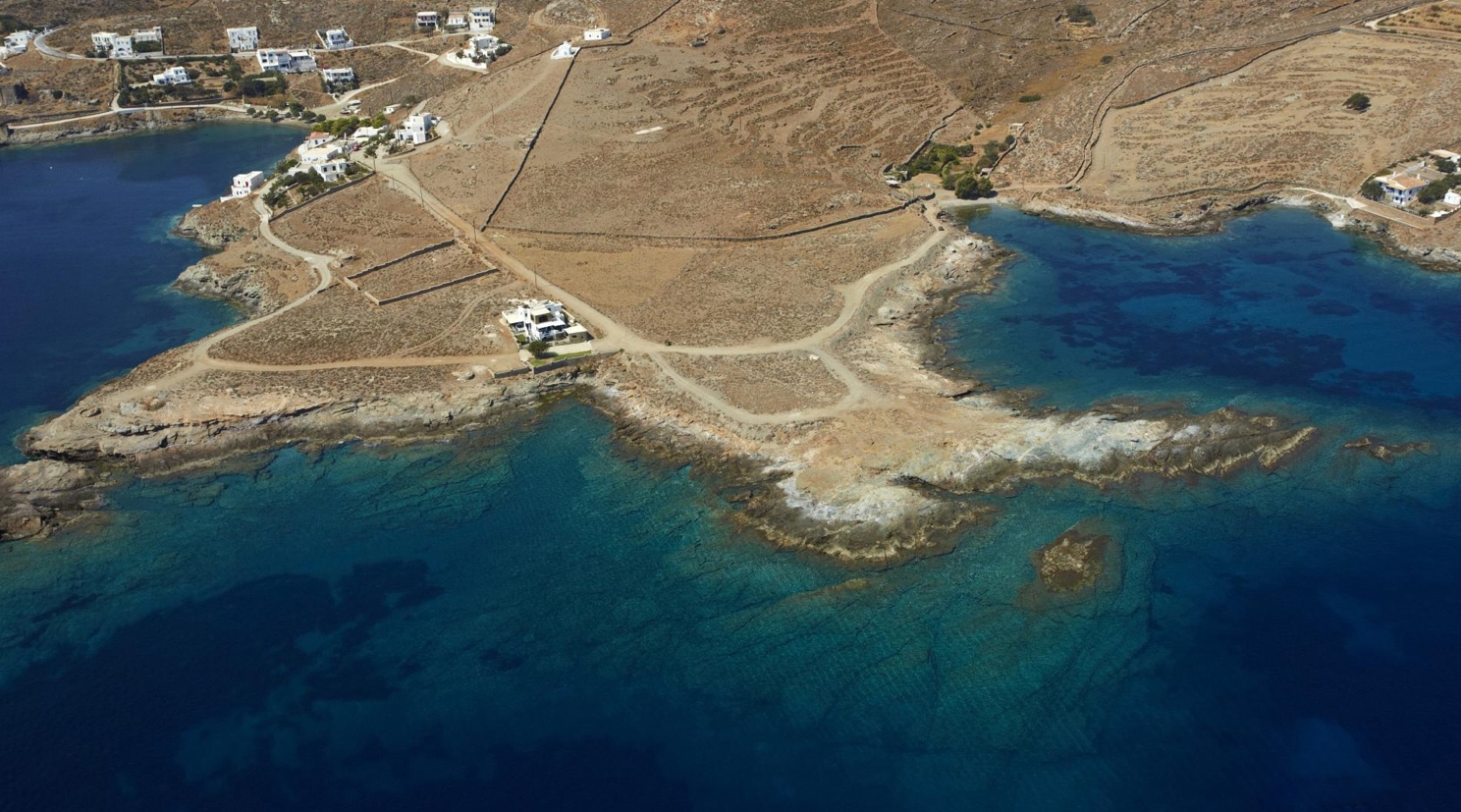

Fig. S10. Aerial photo showing the position of the archeological site of Maroulas on the island of Kythnos.

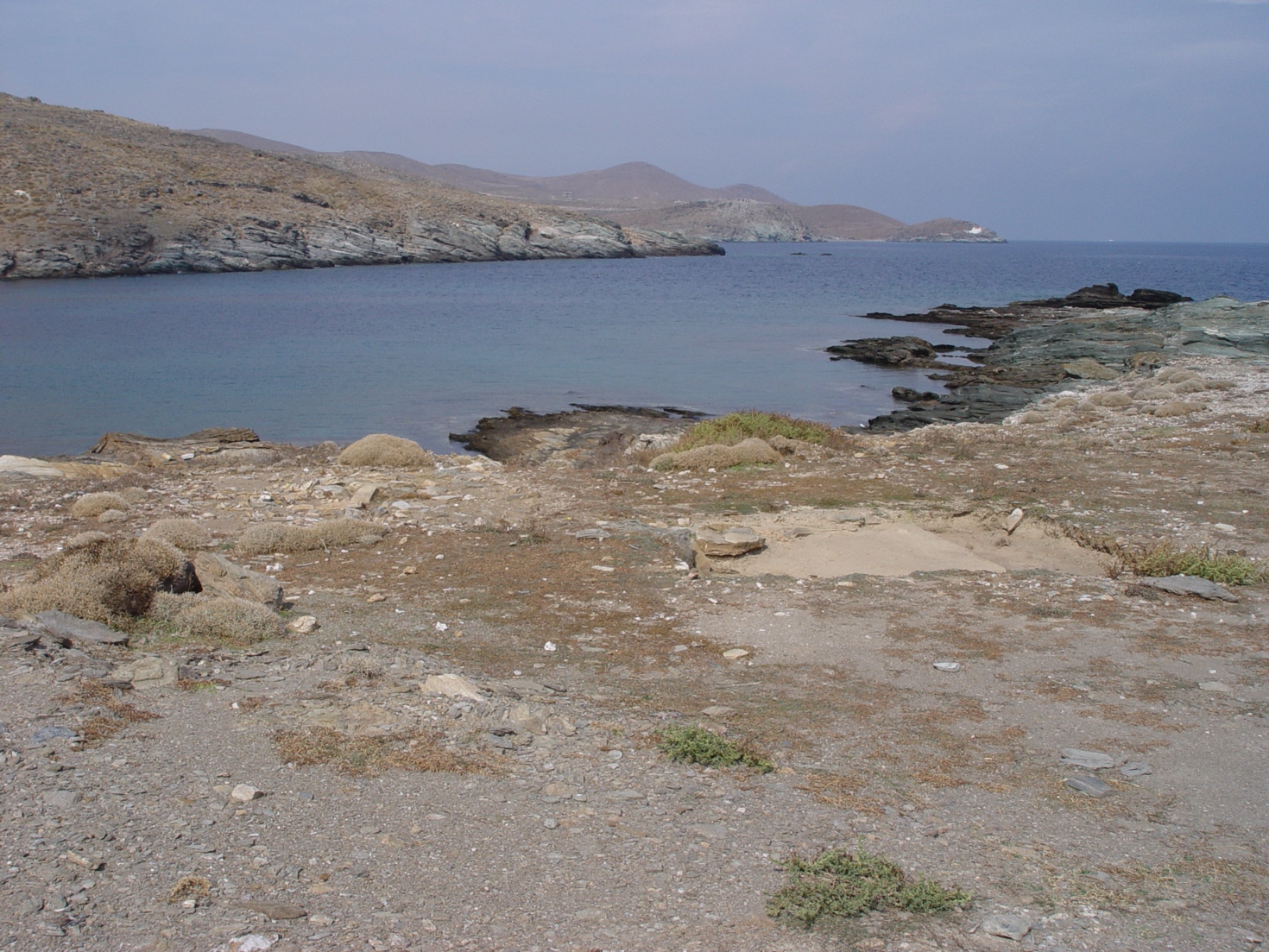

Fig. S11. Position of the archeological site of Maroulas on the island of Kythnos.

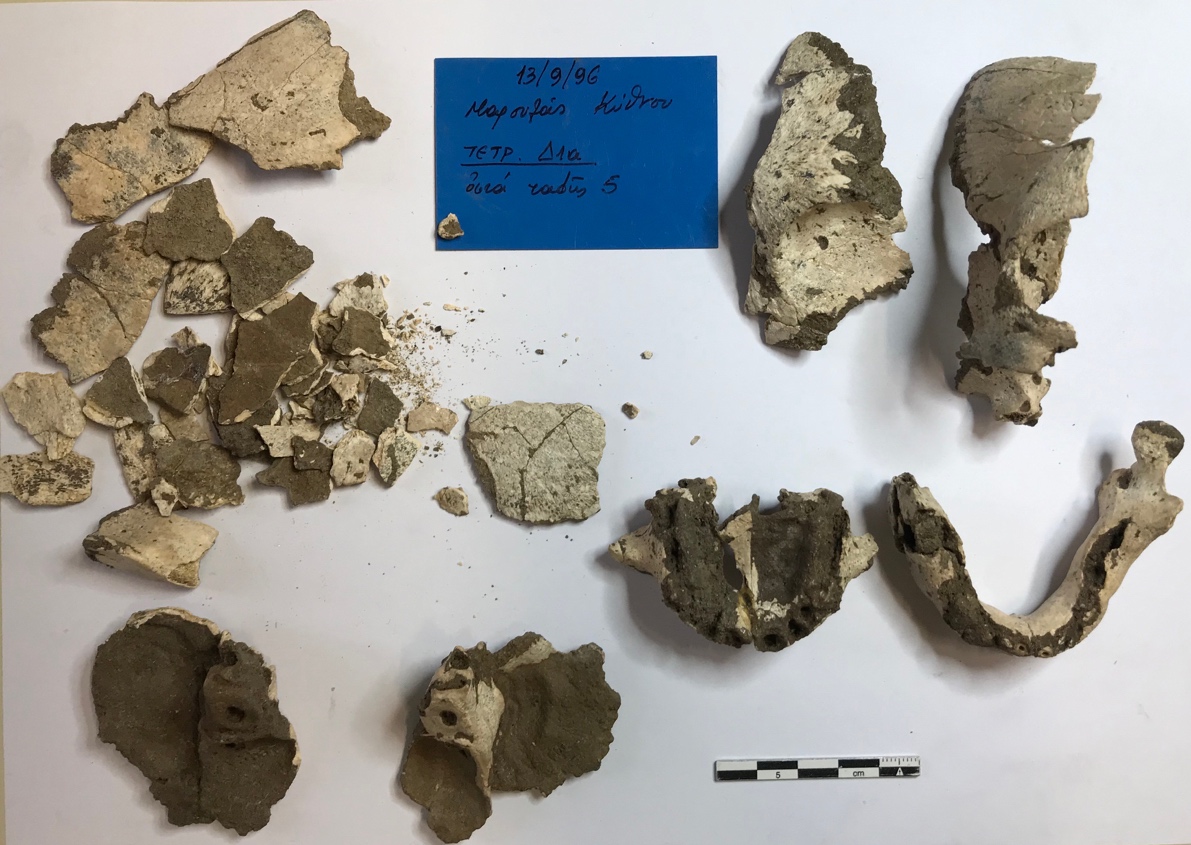

Fig. S12. Sampled human remains marked as Tetr. Δ1a, 5 (13/09/1996, Maroulas-Kythnos).

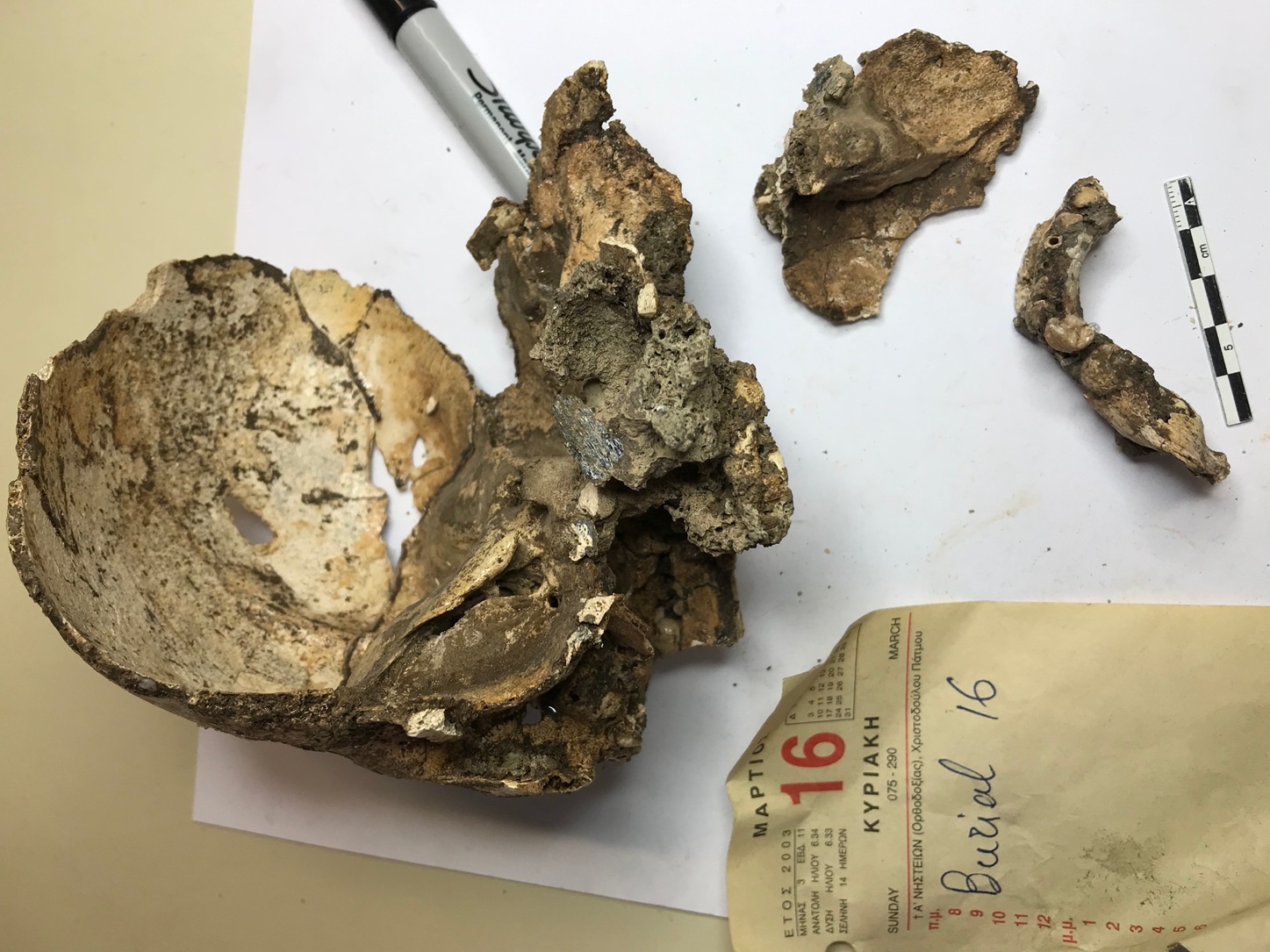

Fig. S13. Sampled human remains marked as Burial 16 (Maroulas-Kythnos).

**S’omu e S’Orku (Sardinia, Italy)**

S’Omu e S’Orku (“The House of the Ogre” in the Sardinian language; hereafter SOMK) (39° 34′ 5″ N, 8° 27′ 36″ E) is a collapsed rockshelter at the base of an aeolianite cliff along the southwestern coast of Sardinia (Fig. S14a). The area is characterized by Quaternary dunes and slope deposits overlying the Paleozoic metamorphic basement. Paleozoic reliefs rise to elevations of up to 300 m. Although SOMK today overlooks the seashore, during the early Holocene the coastline lay several kilometers away, and the shelter opened onto a watercourse (Melis et al. 2023). Only partial access to the deposit is available for research as a huge dune developed over the rockshelter during the Holocene (Melis and Mussi 2026).

The geo-archaeological stratigraphic sequence accumulated between 9500–7800 cal BP encompassing the 9.3k and the 8.2k climatic events of the Early Holocene (Melis and Mussi 2026). During those phases of marked climatic instability, wildfires swept the area, and part of the sequence was built by debris flows with displaced archaeological material. The roof of the rockshelter eventually collapsed sealing the deposit (Melis and Mussi 2026).

The first skeleton was noticed eroding from the sequence, heavily covered in ochre and with a skullcap morphology retaining traits in line with those of pre-LGM Europeans (Fig. S14c), while two more partial skeletons were later discovered in various levels (Melis and Mussi 2016; Oxilia et al. 2025). A fourth burial, from the lowermost part of the deposit, consists of bone fragments and sandy natural casts. Fifty-nine sparse human remains were found incorporated in the debris flows (Melis and Mussi 2026). One of them is an isolated tooth labelled SOMK18/106UM which provided the genetic data presented here (Fig. S15). It was recovered at the base of the debris flow deposits, just above the earliest undisturbed archaeological level (Figure S14b)). This tooth was directly dated to 9396–9030 cal BP at 95% confidence (OxA-42320: 8226 ± 27 BP). The stable isotope values, δ^13^C = –19.6‰ and δ^15^N = 11.1‰, do not suggest a marine reservoir effect for this sample.

The archaeological remains notably include a *Charonia lampas* or trumpet shell with two of the burials (Cristiani et al. 2021), ochre fragments sourced about 50 km south of SOMK (Pisu et al. 2024), and 170 flaked implements, most of them obsidian debris. In the Early Holocene, the mammalian fauna of Sardinia was limited to a few endemic species, represented at SOMK by abundant remains of *Prolagus sardus*, an extinct ochotonid.

Sardinia is an island at a considerable distance from the Italian peninsula and from the European continent and remained such even during the LGM when it formed a large landmass with Corsica (Fig. S14d). The scarce archaeological evidence of early Holocene age from both Sardinia and nearby Corsica, together with the environmental instability recorded at SOMK, suggest a limited Mesolithic peopling which became extinct some centuries before Neolithic groups settled the islands and introduced new plant and animal species.

*References:* Cristiani et al. 2021; Melis et al. 2023; Melis and Mussi 2016; Melis and Mussi 2026; Oxilia et al. 2025; Pisu et al. 2024.

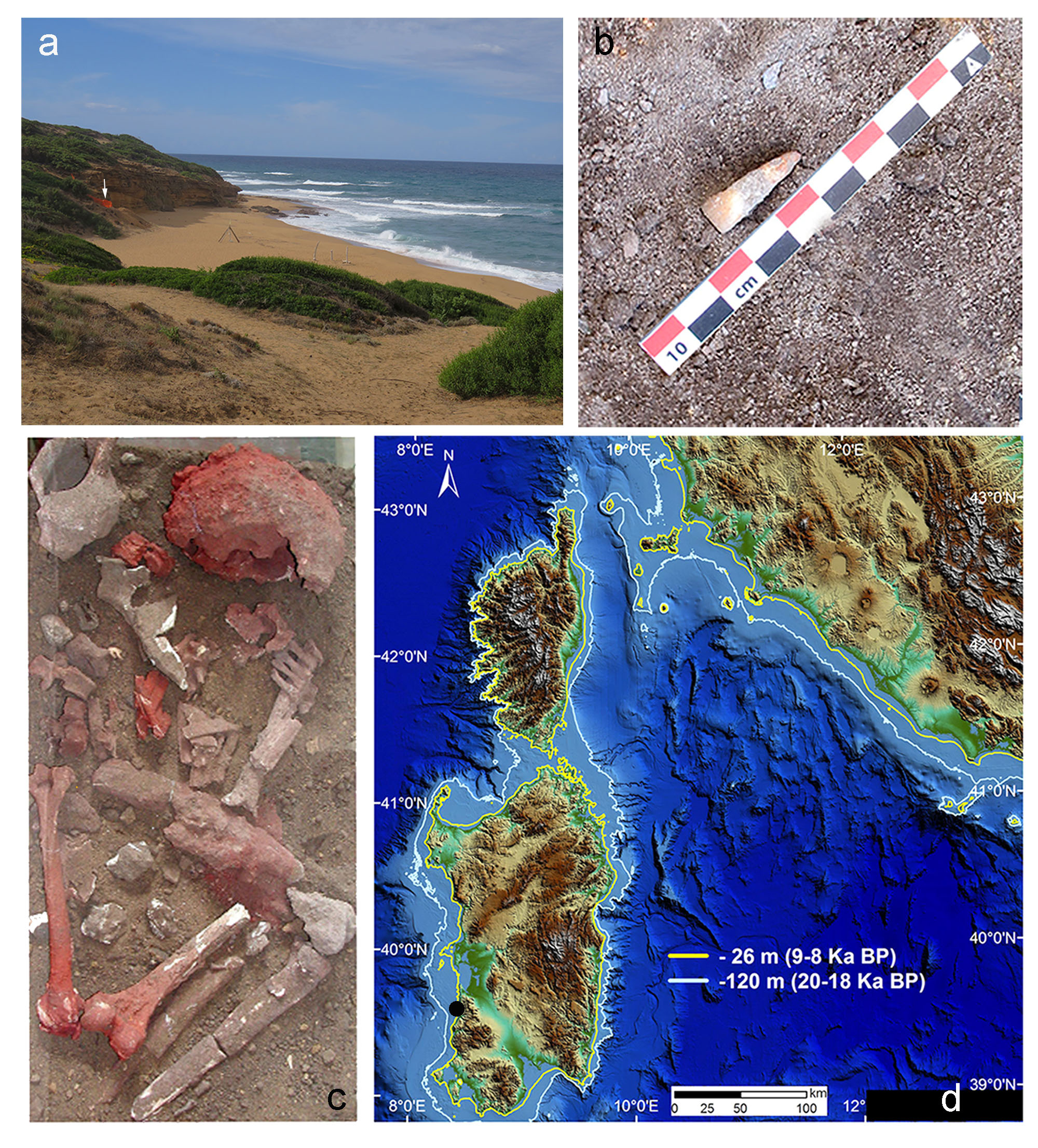
 Figure S14. The S’Omu e S'Orku (SOMK) site. (a) View of the coastal area with the SOMK site (white arrow) on the south-western coast of Sardinia (Italy); (b) the human tooth labelled SOMK18/106UM in stratigraphic context, lying at the base of the debris flow deposits and just above the earliest undisturbed archaeological level; (c) the skeletal remains labelled SOMK1 heavily covered by ochre; (d) the palaeocoastline at –26 m (9–8 ka BP Mesolithic times) and –120m (20–18 ka BP LGM) along Sardinia, Corse, and Elba. The black dot indicates the SOMK site.

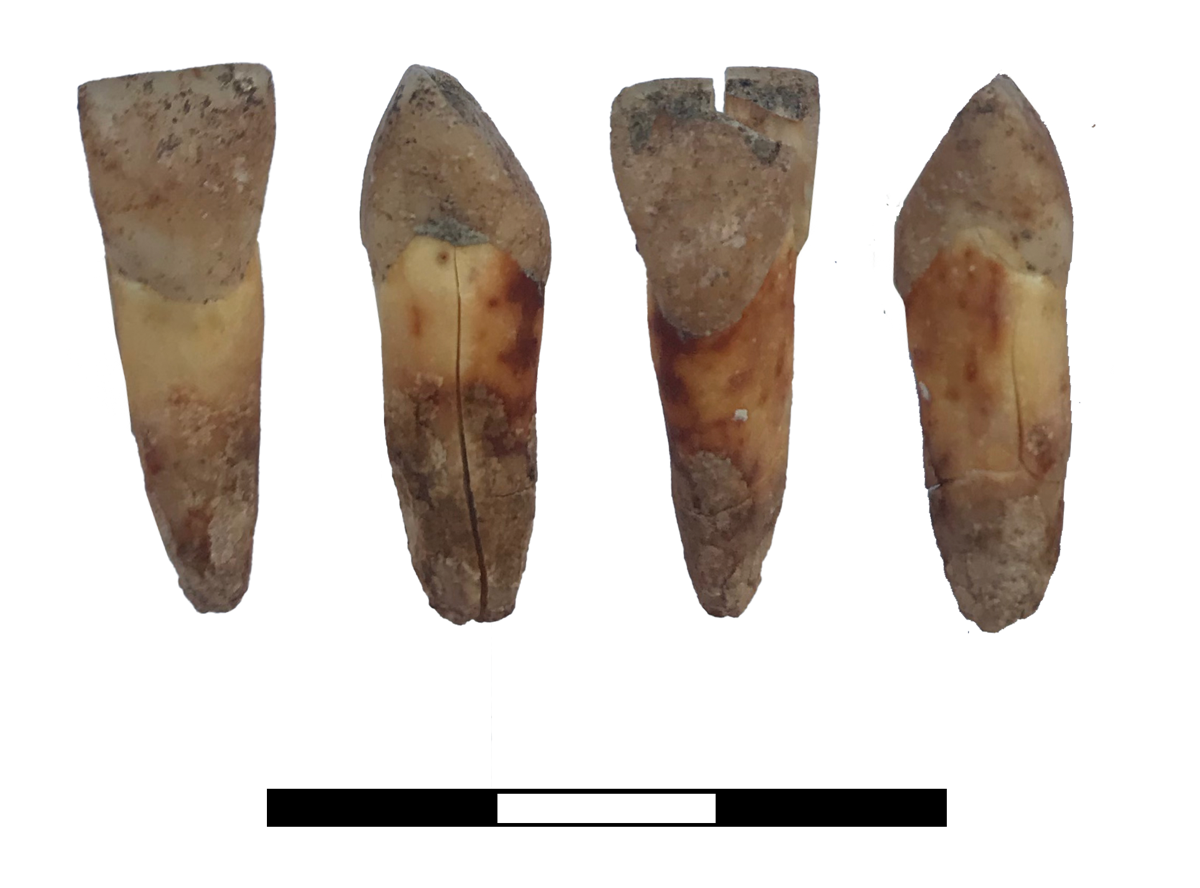

Fig. S15. Sampled human tooth labelled as SOMK18/106UM.

**Romanelli (Apulia, Italy)**

Grotta Romanelli (40° 00′ 58.30″ N, 18° 26′ 00.01″ E), is located on the southernmost eastern coast of Apulia, Italy, at 7.3 m above present-day sea level (Fig. S16). The cave is considered the first Upper Palaeolithic site discovered in Italy, with investigations commencing in 1900 (Blanc 1928). The sequence consists of deposits dating to the Late Pleistocene. The stratigraphy is divided into a lower complex encompassing the beach deposit (level K), the bone breccia (level I), the stalagmitic layer (level L), and the *terre rosse* (red soils, level G), dating to MIS 9–7 (based on new U/Th dates; Pieruccini et al. 2022) and MIS 5e, and an upper complex comprising the *terre brune* (brown soils, levels E–A), dating to the Upper Palaeolithic and Mesolithic. A stalagmitic layer (level F) separates the two complexes (Fabbri et al. 2003). A more recent phase of research at the site began in 2015 and focused on the upper parts of the sequence, spanning the Epigravettian to the Mesolithic. New radiocarbon dates suggest that level D encompasses the Late Palaeolithic between c. 14000 and 12500 cal BP; level C, overlying level D, dates to between c. 12500 and 10700 cal BP; and level B dates to between c. 9500 and 8600 cal BP (Calcagnile et al. 2019).

Faunal remains have been collected throughout the sequence, especially from the *terre rosse* and the *terre brune*. The assemblage from these latter levels is extremely large (Tagliacozzo 2003) and documents human exploitation of mammals, in particular red deer, aurochs, and *Equus hydruntinus* (Fiore et al. 2003), but also small taxa (e.g., fox, wildcat, hare, badger, and even hedgehog) as indicated by butchery marks, types of bone fractures, and localized burning (Tagliacozzo and Fiore 1998). Birds represented an important supplement to human diet (Cassoli and Tagliacozzo 1997), but fish was occasionally exploited as well. The site yielded some of the earliest evidence for the presence of dogs in Italy (Boschin et al. 2020).

Examples of parietal art as well as portable engraved objects have been found at the site (Sigari 2026; Sigari et al. 2021). Human remains recovered from the cave include three articulated burials (one adult and two children) found in the *terre brune*, as well as various disarticulated human remains (cranial and postcranial elements, mandibles, and isolated teeth) (Fabbri 1987; Fedele 2003; Mecozzi et al. 2022). While some human remains have been attributed to levels C and D, others lack more precise contextual information.

The human rib fragment (ZooMS R16) from which we obtained genetic data derives from layer V, which corresponds to the upper part of the level C of the main stratigraphy and was excavated in 1958 (Fig. S17). It was identified within the bone collection through a systematic search for human remains among the less identifiable bone fragments stored at the Museo delle Civiltà in Rome, Italy. We applied collagen peptide fingerprinting, known as Zooarchaeology by Mass Spectrometry (ZooMS), to identify the fragment as human (Fig. S18). The ZooMS analysis followed the protocol published by Buckley et al. (2009; see also van der Sluis et al. 2014). Briefly, bone samples were demineralized overnight in 0.6 M hydrochloric acid, followed by gelatinisation of the acid-insoluble residue in 50 mM ammonium bicarbonate at 65°C. The resulting supernatant was digested overnight at 37°C with sequencing-grade trypsin (Promega, UK) and subsequently acidified with 5% trifluoroacetic acid. Next, 0.5 μL of the sample solution was co-crystallised with 0.5 μL of saturated α-cyano-4-hydroxycinnamic acid matrix solution on a Bruker ground steel matrix-assisted laser desorption/ionisation time-of-flight (MALDI-TOF) target plate. Samples were analysed using a Bruker UltrafleXtreme mass spectrometer equipped with a frequency-tripled Nd:YAG laser at the Department of Chemistry, Columbia University. A total of 2000 laser shots were acquired over the m/z range 800–3700. Spectra were manually screened for taxonomically informative markers by comparing them with the reference datasets published by Buckley (2016), Buckley et al. (2009, 2017), and Welker et al. (2015).

The AMS radiocarbon date obtained for this fragment calibrates to 13225–12765 cal BP at 95% confidence (OxA-X-3144-12: 11110 ± 110 BP), fits with its provenience from level C and was accompanied by the following stable isotope values measured as indicative AMS burns: δ^13^C = –19.7‰ and δ^15^N = 15.2‰. These values do not suggest the presence of a marine reservoir effect but instead indicate a diet with a high contribution of terrestrially derived protein.

*References:* Blanc 1928; Boschin et al. 2020, Calcagnile et al. 2019; Cassoli, Tagliacozzo 1997; Fabbri 1987; Fabbri et al. 2003; Fedele 2003; Fiore et al. 2003, Mecozzi et al. 2022; Pieruccini et al. 2022; Sigari 2026; Sigari et al. 2021; Tagliacozzo 2003; Tagliacozzo, Fiore 1998.

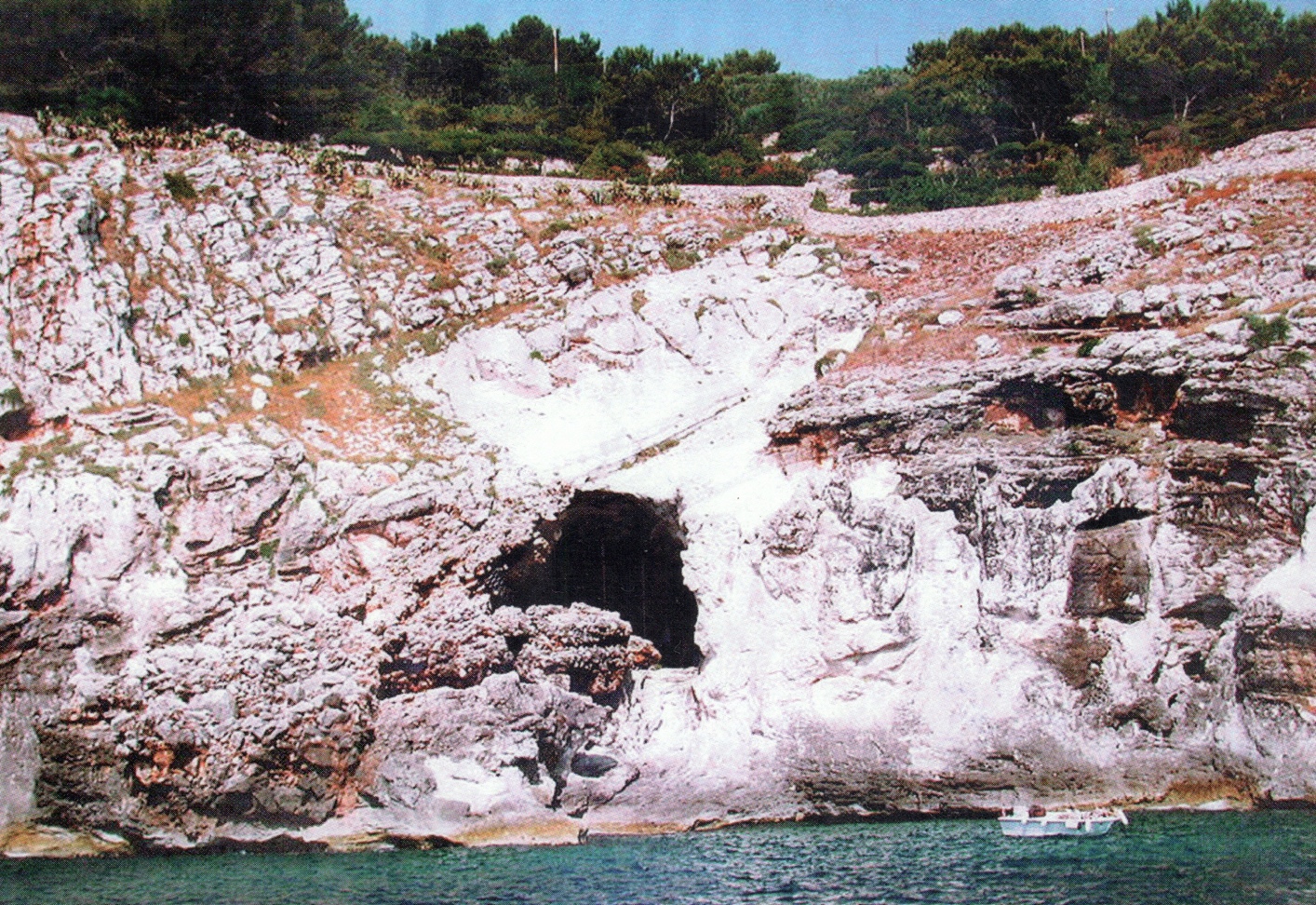

Fig. S16. The site of Grotta Romanelli from the sea.

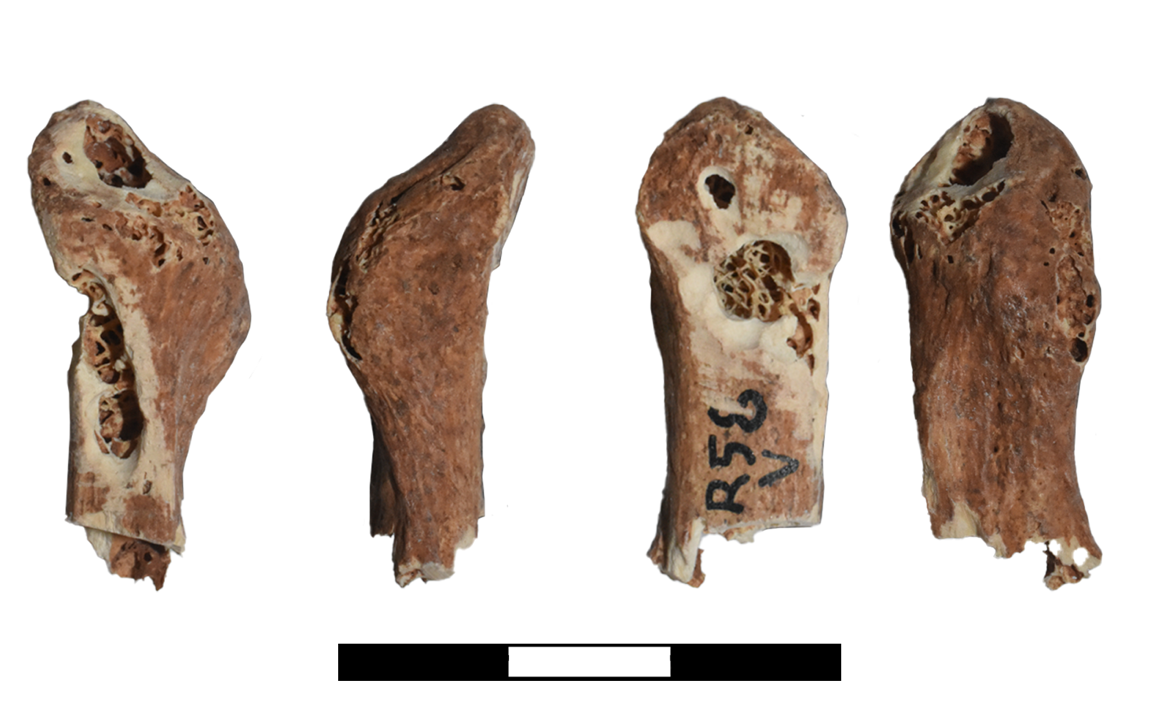

Fig. S17. Sampled rib from Romanelli (R58, V, ZooMS R16).

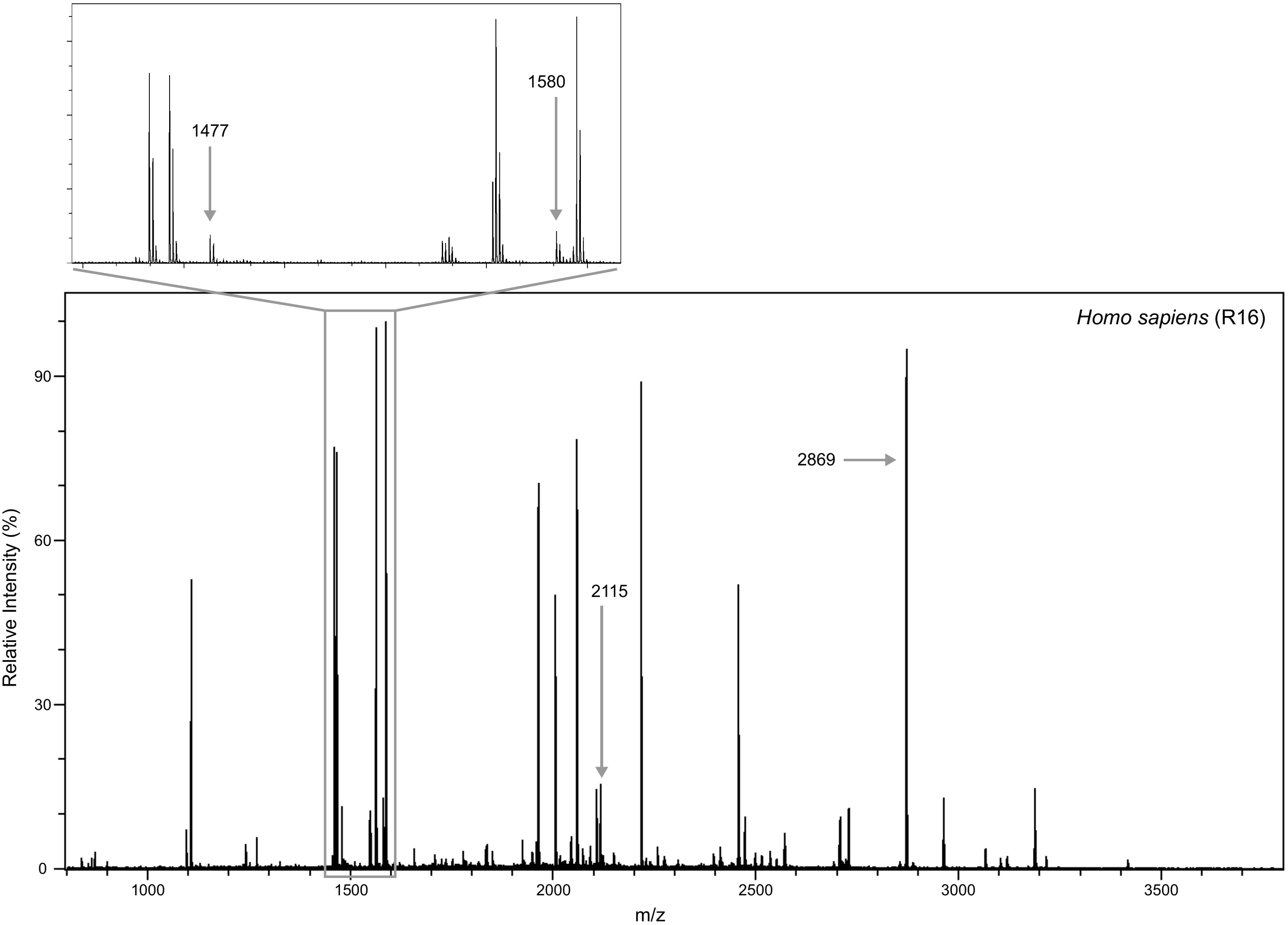

Fig. S18. MALDI spectra identification of the rib fragment from Romanelli. Masses of the key markers used for taxonomic identification as Homo are indicated with arrows (m/z 1477, 1580, 2115, 2869).

Borić, D., Cristiani, E., Zupancich, A., Mecozzi, B., Fasser, N., Vinet, A., Fontana, F., Hajdinjak, M., Pelin Sümer, A., Monaco, L., Lepić, A., Marić, A., Karaica, A., & Whallon, R. Badanj Revisited: An Epigravettian Persistent Place in the Adriatic Hinterland of the Dinaric Alps. *PLOSOne* (forthcoming).

Boroneanţ, A. The Mesolithic in Banat, in N. Tasić & F. Draşovean (eds.) *The Prehistory of Banat. I: The Palaeolithic and Mesolithic*. Bucharest, The Publishing House of the Romanian Academy, 103–141 (2011).

Boroneanţ, V. La période épipaléolithique sur la rive roumaine des Portes de Fer du Danube. *Praehistorische Zeitschrift* **45(1)**, 1–25 (1970).

Boroneanţ, V. *Paleolithique superieur et epipaleolithique dans la zone des Portes de Fer*. Bucureşti, Silex (2001).

Boschin, F., Bernardini, F., Pilli, E., Vai, S., Zanolli, C., Tagliacozzo, A., Fico, R., Fedi, M., Corny, J., Derossi, D., Lati, M., Modi, A., Vergata, C., Tuniz, C., Moroni, A., Boscato, P., Caramelli, D. & Ronchitelli, A. The First Evidence for Late Pleistocene Dogs in Italy. *Scientific Reports* **10**, 13313 (2020) <https://doi.org/10.1038/s41598-020-69940-w>

Buckley, M., Collins, M., Thomaes‐Oates, J. *et al.* Species identification by analysis of bone collagen using matrix‐assisted laser desorption/ionisation time‐of‐flight mass spectrometry. *Rapid Communications in Mass Spectrometry* **23**, 3843–3854 (2009).

Buckley, M., Harvey, V. L., & Chamberlain, A. T. Species identification and decay assessment of Late Pleistocene fragmentary vertebrate remains from Pin Hole Cave (Creswell Crags, UK) using collagen fingerprinting. *Boreas* **46(3)**, 402–411 (2017).

Buckley, M. Species identification of bovine, ovine and porcine type 1 collagen; comparing peptide mass fingerprinting and LCbased proteomics methods. *International Journal of Molecular Sciences* **17(4)**, 445 (2016).

Cassoli, P.F. & Tagliacozzo, A. Butchering and cooking of birds in the palaeolithic site of Grotta Romanelli (Italy). *International Journal of Osteoarchaeology* **7**, 303–320 (1997).

Cook, G.T., Bonsall, C., Hedges, R.E.M., McSweeney, K., Boroneanț, V., Bartosiewicz, L., & Pettitt, P.B. Problems of Dating Human Bones from the Iron Gates. *Antiquity* **76**, 77–85 (2002).

Cristiani, E., R. T. Melis, & M. Mussi, Marine Shells as Grave Goods at S’Omu e S’Orku (Sardinia, Italy), in D. Borić, D. Antonović, & B. Mihailović (eds.) *Foraging Assemblages*, Belgrade and New York: Serbian Archaeological Society and the Italian Academy for Advanced Studies in America, Columbia University, 558–566 (2021).

Cullen, T. Mesolithic Mortuary ritual at Franchthi Cave, Greece. *Antiquity* **69(263)**, 270–289 (1995).

Cullen, T. with the coll. of A. Papathanasiou, *Funerary rituals and human biology at Franchthi,* Indiana University Press, Bloomington (forthcoming).

Calcagnile, L., Sardella, R., Mazzini, I., Giustini, F., Brilli, M., D’Elia, M., Braione, E., Conti, J., Mecozzi, B., Bona, F., Iurino, D.A., Lembo, G., Muttillo, B., & Quarta, G. New Radiocarbon Dating Results from The Upper Paleolithic–Mesolithic Levels, Grotta Romanelli (Apulia, Southern Italy). *Radiocarbon* **61(5)**, 1211–1220 (2019).

Fabbri, P. F. Restes humains retrouvés dans la grotte Romanelli (Lecce, Italie): étude anthropologique. *Bull Mém Soc Anthropol Paris* **4(4)**, 219–247 (1987).

Fabbri, P. F., Ingravallo, E., & Mangia, A. (eds.). *Grotta Romanelli nel centenario della sua scoperta (1900–2000)*, Lecce, Congedo Editore (2003).

Facorellis, Y., Damiata, B. N., Vardala-Theodorou, E., Ntinou, M. & Southon, J. AMS Radiocarbon Dating of The Mesolithic Site Maroulas on Kythnos and Calculation of the Regional Marine Reservoir Effect, in A. Sampson, M. Kaczanowska, & J.K. Kozlowski (eds.) *The Prehistory of the Island of Kythnos (Cyclades, Greece) and the Mesolithic Settlement at Maroulas*, Krakow, The Polish Academy of Arts and Sciences and the University of the Aegaen–Rhodes, 127–135 (2010).

Fedele, F. L’acquisto della collezione Stasi da parte dell’Università di Napoli, in Fabbri, P. F., Ingravallo, E., & Mangia, A. (eds.) *Grotta Romanelli nel centenario della sua scoperta (1900–2000)*, Lecce, Congedo Editore, 27–38 (2003).

Fiore I., Curci A., & Tagliacozzo A. Tecniche di macellazione e sfruttamento dei grandi ungulati (*Bos primigenius, Equus hydruntinus, Cervus elaphus*) dei livelli epigravettiani di Grotta Romanelli (scavi 1954 e 1958), in Fabbri, P. F, Ingravallo, E. & Mangia, A. (eds.), Grotta Romanelli nel centenario della sua scoperta 1900–2000, Lecce, Congedo Editore, 149–168 (2003).

Jakobsen, T. W. Excavations at Porto Cheli and vicinity, preliminary report, II: The Franchti Cave. *Hesperia* **38**, 343–381 (1969).

Jakobsen, T. W. 17,000 Years of Greek Prehistory, in *Hunters, Farmers and Civilizations: Old World Archaeology*. *Readings from Scientific American*, San Francisco, W. H. Freeman and Company, 133–151 (1976).

Martinoia, V., Papathanasiou, A., Talamo, S., MacDonald, R. & Richards, M. P. High-Resolution Isotope Dietary Analysis of Mesolithic and Neolithic Humans from Franchthi Cave, Greece, *PloSOne* (2025) <https://doi.org/10.1371/journal.pone.0310834>

Mathieson, I., *et al.* The Genomic History of Southeastern Europe. *Nature* **55**, 197–203 (2018). <https://doi.org/10.1038/nature25778>

Mecozzi, B., Buzi, C., Iannucci, A., Micarelli, I., Bona, F., Forti, L., Lembo, G., Manzi, G., Mazzini, I., Muttillo, B., Pieruccini, P., Ranaldo, F., Sigari, D., & Sardella, R. 2022. New Human Fossil from the Latest Pleistocene Levels of Grotta Romanelli (Apulia, Southern Italy). *Archaeological and Anthropological Sciences* **14:27** (2022) <https://doi.org/10.1007/s12520-021-01491-1>

Melis R.T., Demurtas V., Mussi M., Orrù P.E., Sulis A., Altamura F., Erbì R., Orrù M., & Deiana G. The paleolandscape evolution of the southwestern coast of Sardinia (Italy) and its impact on Mesolithic settlements.  *Journal of Maps* 19 (2023) 2182722 doi.org/[10.1080/17445647.2023.2182722](https://doi.org/10.1080/17445647.2023.2182722)

Melis, R. T., Mussi, M., Floris, R., Lamothe, M., Palombo, M. R., & Usai A. Popolamento e ambiente nella Sardegna centro occidentale durante l’Olocene antico: primi risultati. In *Atti della XLIV Riunione Scientifica dell’Istituto Italiano di Preistoria e Protostoria: La Preistoria e la Protostoria della Sardegna: Cagliari, Barumini, Sassari 23–28 novembre 2009*, 427–434. Firenze (2012).

Melis, R. T. & Mussi, M. Mesolithic burials at S’Omu e S’Orku (SOMK) on the south-western coast of Sardinia, in J. M. Grünberg, B. Gramsch, L. Larsson, J. Orschiedt, and H. Meller (eds.) *Mesolithic Burials – Rites, Symbols and Social Organisation of Early Postglacial Communities. International Conference Halle (Saale), Germany, 18th–21st September 2013 (Tagungen des Landesmuseums für Vorgeschichte Halle 13/II)*, 733–40. Halle (Saale), Landesamt für Denkmalpflege und Arch.ologie Sachsen-Anhalt, Landesmuseum für Vorgeschichte (2016).

Melis R.T. & Mussi M. The Mesolithic seen through a keyhole at S’Omu e S’Orku (SOMK) and the case of environmental hazards on the western coast of Sardinia, 9,500–7,800 cal BP. *The Holocene* (2026) <https://doi.org/10.1177/09596836261432439>

Munro, N. D. & Stiner, M. C. Zooarchaeological Evidence for Early Neolithic Colonization at Franchthi Cave (Peloponnese, Greece). *Current Anthropology* **56**, 596–603 (2015). <https://doi.org/10.1086/682326>

Oxilia G., Mussi M., Chiriu D., Pisu F.A., Marini E., & Melis R.T. Virtual Analysis of a Concretioned Skullcap from S’Omu e S’Orku, an Early Holocene Mesolithic site of Sardinia. *American Journal of Biological Anthropology* **187**, e70065 (2025) [doi.org/10.1002/ajpa.70065](https://doi.org/10.1002/ajpa.70065)

Ottoni, C., Borić, D., Cheronet, O., Sparacello, V., Dori, I., Coppa, A., Antonović, D., Vujević, D., Price, T.D., Pinhasi, R., & Cristiani, E. Tracking the Transition to Agriculture in Southern Europe Through Ancient DNA Analysis of Dental Calculus. *Proceedings of the National Academy of Sciences of the USA* **118(32)**, e2102116118 (2021).

Perlès, C. Long-Term Perspectives in the Occupation of the Franchthi Cave: Continuity and Discontinuity, in G. N. Bailey, E. Adam, E. Panagopoulou, C. Perlès, & K. Zachos (eds.) *The Palaeolithic Archaeology of Greece and Adjacent Areas*. Athens, British School at Athens Studies 3, 311–318 (1999).

Perlès, C. *The Early Neolithic in Greece. The First Farming Communities in Europe*, Cambridge, Cambridge University Press (2001).

Perlès, C. *Ornaments and other ambiguous artifacts. Vol. 2. The Neolithic*. Bloomington, Indiana University Press (2023).

Perlès, C. & Rose, M. *35,000 years of occupation at Franchthi Cave: A Chronostratigraphic Synthesis*, Bloomington, Indiana University Press (forthcoming).

Pieruccini, P., Forti, L., Mecozzi, B., Iannucci, A., Yu, T.‑L., Shen, Ch.‑Ch., Bona, F., Lembo, G., Muttillo, B., Sardella, R., & Mazzini, I. Stratigraphic reassessment of Grotta Romanelli sheds light on Middle‑Late Pleistocene palaeoenvironments and human settling in the Mediterranean. *Scientific Reports* **12**, 13530 (2022). <https://doi.org/10.1038/s41598-022-16906-9>

Pisu F., Porcu S., Carboni R., Mameli V., Cannas C. Naitza S., Melis R., Mussi M., Chiriu D. Innovative method for provenance studies in Cultural Heritage: A New Algorithm Based on Observables from High-Resolution Raman Spectra. *Spectrochimica Acta Part A: Molecular and Biomolecular Spectroscopy* **329**, 125581 (2024). doi.org/10.1016/j.saa.2024.125581

van der Sluis, L. G. *et al*. Combining histology, stable isotope analysis and ZooMS collagen fingerprinting to investigate the taphonomic history and dietary behaviour of extinct giant tortoises from the Mare aux Songes deposit on Mauritius. *Palaeogeography, Palaeoclimatology, Palaeoecology* **416**, 80–91 (2014).

Sampson, A. 2010. *Τhe Prehistory of the Island of Kythnos and the Mesolithic Settlement at Maroulas*. Krakow: Polish Academy of Sciences and Arts and University of the Aegean.

Sigari, D. The Figurative Motifs in the Portable Art of Grotta Romanelli (Southern Italy) Within the Late Pleistocene Art Tradition of Southwestern Europe. *Journal of Archaeological Science: Reports* **71**, 105630 (2026).

Sigari, D., Mazzini, I., Conti, J., Forti, L., Lembo, G., Mecozzi, B., Muttillo, B., & Sardella, R. Birds and bovids: new parietal engravings at the Romanelli Cave, Apulia. *Antiquity* **95(384)**, 1387–1404 (2021).

Tagliacozzo, A. Archeozoologia dei livelli dell’Epigravettiano finale di Grotta Romanelli (Castro, Lecce) Strategie di caccia ed economia di sussistenza, in Fabbri P. F, Ingravallo E. & Mangia A. (eds.) *Grotta Romanelli nel centenario della sua scoperta 1900-2000*, Lecce, Congedo Editore, 169–216 (2003).

Tagliacozzo, A. & Fiore, I. Butchering of Small Mammals in the Epigravettian Levels of the Romanelli Cave (Apulia, Italy), in Actes du XVIII Rencontre Internationale d'archéologie et d'histoire d'Antibes, *Economie Préhistoriques*, Editions APDCA, Sophia Antipolis, 413–423 (1998).

Vitelli, K. D. *Franchthi Neolithic Pottery. Vol. 1: Classification and Ceramic Phases 1 and 2*, Bloomington, Indiana University Press (1993).

Welker, F., Soressi, M., Rendu, W. *et al.* Using ZooMS to identify fragmentary bone from the late Middle/Early Upper Palaeolithic sequence of Les Cottés, France. *Journal of Archaeological Science* **54**, 279–286 (2015).

Whallon, R. *Badanj: Epipalaeolithic Excavations in Herzegovina, 1986–1987*, Ann Arbor, University of Michigan Museum of Anthropological Archaeology, Memoirs 67 (2025).
